# Spurious regulatory connections dictate the expression-fitness landscape of translation termination factors

**DOI:** 10.1101/2020.11.27.400200

**Authors:** Jean-Benoît Lalanne, Darren J. Parker, Gene-Wei Li

## Abstract

During steady-state cell growth, individual enzymatic fluxes can be directly inferred from growth rate by mass conservation, but the inverse problem remains unsolved. Perturbing the flux and expression of a single enzyme could have pleiotropic effects that may or may not dominate the impact on cell fitness. Here we quantitatively dissect the molecular and global responses to varied expression of translation termination factors (peptide release factors, RFs) in bacterium *Bacillus subtilis*. While endogenous RF expression maximizes proliferation, deviations in expression lead to unexpected distal regulatory responses that dictate fitness reduction. Molecularly, RF depletion causes expression imbalance at specific operons, which activates master regulators and detrimentally overrides the transcriptome. Through these spurious connections, RF abundances are thus entrenched by focal points within the regulatory network, in one case located at a single stop codon. Such regulatory entrenchment suggests that predictive bottom-up models of expression-fitness landscapes will require near-exhaustive characterization of parts.

**Highlights:**

- Precision measurements enable multiscale expression-to-fitness mapping.
- RF depletion leads to imbalanced translation for co-transcribed gene pairs.
- Imbalanced translation induces unintended regulons to the detriment of cell fitness.
- Swapping a single stop codon rewires global susceptibility to RF perturbation.

## Introduction

Formulating predictive models connecting genetic information to phenotypes constitutes an overarching goal in genomics and systems biology (Lopatkin and Collins, 2020; Ostrov et al., 2019; Shendure et al., 2019). In single-celled microbes, the relationship between genotype and phenotype can be conceptually decomposed into two distinct maps; the first relating genome sequence to gene expression, and the second, connecting gene expression to whole-cell properties such as proliferation. Rapid progress on the characterization of *cis*-regulatory elements, spurred by integration of massively parallel reporter assays (Patwardhan et al., 2009, 2012; Sharon et al., 2012) with novel computational frameworks (de Boer et al., 2020; Bogard et al., 2019; Jaganathan et al., 2019; Rosenberg et al., 2015), has achieved headway in predicting expression features from DNA sequence (Agarwal and Shendure, 2018; Cambray et al., 2018; Sample et al., 2019; Urtecho et al., 2020). By contrast, expression-fitness landscapes, defined as the distinct relationships between the expression level of individual genes and the cell fitness, remain understudied despite being the basis of selective pressures on protein abundances. This information gap limits both the engineering of complex biological functions and the interpretability of genetic variation.

Predicting the shape of expression-fitness landscapes requires quantitative characterization of the cellular state at multiple levels. Although numerous expression-fitness landscapes have been previously mapped (Chou et al., 2014; Dekel and Alon, 2005; Duveau et al., 2017; Hawkins et al., 2019; Jost et al., 2020; Kavčič et al., 2019; Keren et al., 2016; Knöppel et al., 2016; Li et al., 2014; Palmer et al., 2018; Parker et al., 2020; Schober et al., 2019; Tubulekas and Hughes, 1993), these measurements rarely include concomitant assessment of the internal cell state following perturbations (but see (Jost et al., 2020; Parker et al., 2020)). With limited information bridging the molecular to cellular scales, the root causes of observed fitness defects are challenging to pinpoint (Fig. 1a). In particular, changes in enzyme levels not only directly affect flux and growth (Fig. 1a, inset i for the case of translation factors) (Ehrenberg and Kurland, 1984; Klumpp et al., 2013; Li et al., 2014), they can also have indirect pleiotropic effects that take the form of damage propagation across three connected levels of biological organization (Fig. 1a, inset ii). First, a reduced enzymatic flux could affect the expression of other genes via specific molecular mechanisms (mechanistic level). Second, these proximal changes in expression could ripple through the regulatory network, leading to further changes in expression genome-wide (regulatory level). Third, each terminal downstream expression change could have an impact on fitness (systemic level). Whether selective pressures on the abundance of proteins predominantly relate to direct impacts on flux or are rather dominated by indirect cellular responses remains unresolved.

**Fig. 1.**
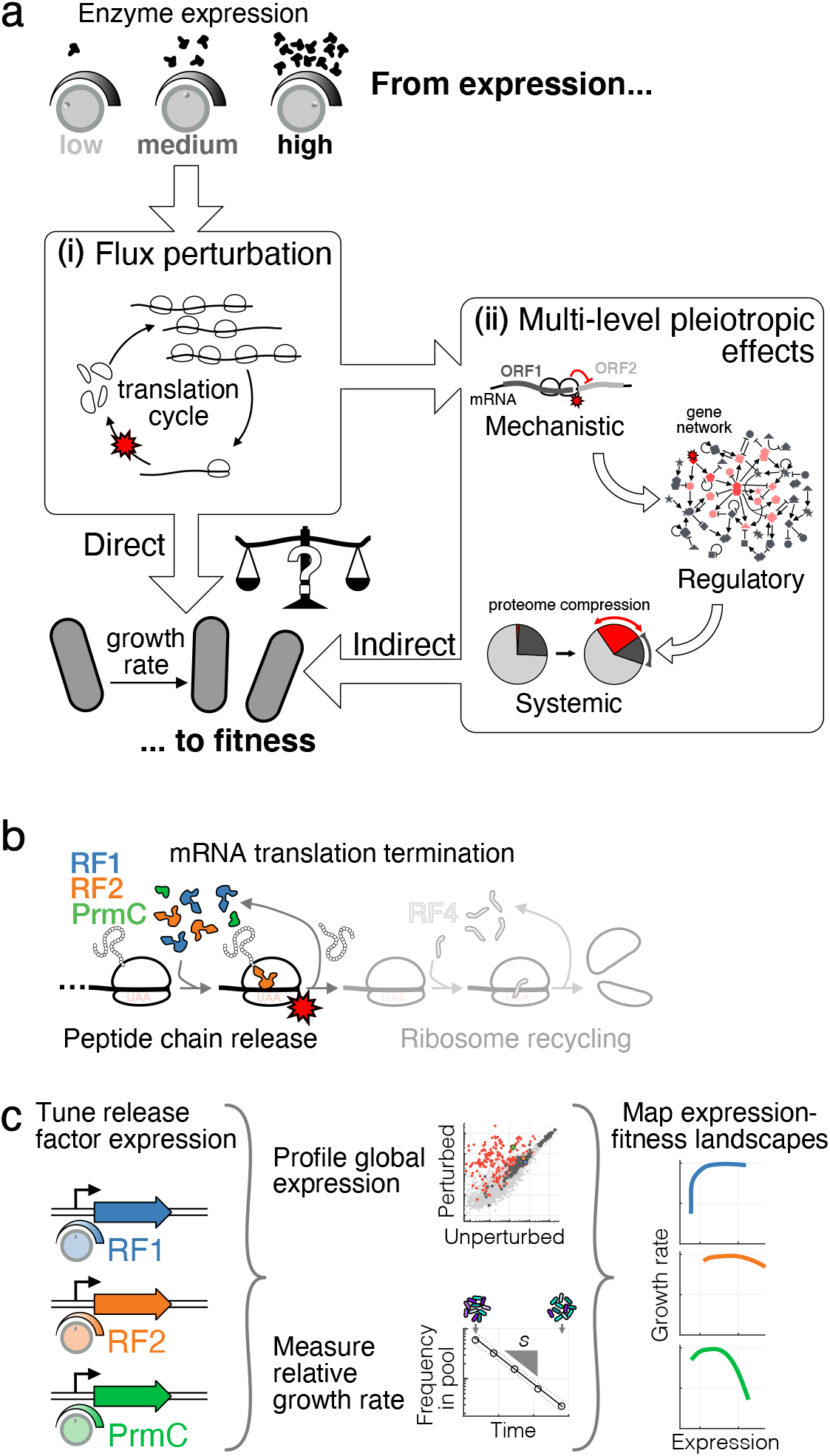
Mapping the underlying determinants of the release factor expression-fitness landscapes. **(a)** Expression-fitness landscapes, which connect enzyme expression (microscopic variable) to the growth rate (cellular phenotype), can be dictated by direct, or indirect effects. Inset (i): direct effects correspond to reduction in the flux cognate to the perturbed enzyme (protein synthesis rate in the case of translation factors). Inset (ii): indirect effects result from pleiotropic propagation across mechanistic, regulatory, and systemic levels. **(b)** As a case study, the expression of peptide chain release factors (RFs: RF1, RF2, and associated methyltransferase PrmC), involved in the first step of mRNA translation termination, was tuned around endogenous levels. **(c)** Strains with inducible copies of RFs, and deleted endogenous genes, were used to systematically vary RF expression. The resulting impacts on the cell internal state (RNA-seq, ribosome profiling) and relative growth rate *s* (competition experiments) were measured, leading to precise mapping of expression-fitness landscapes. See Fig. S1 and S2 for details on strains and measurement platform.

Here as a case study, we systematically vary the expression of enzymes involved in the core process of mRNA translation termination (Fig. 1b) in Gram-positive bacterium *Bacillus subtilis*. We focus on peptide chain release factors, RF1, RF2, and their methyltransferase PrmC (hereafter collectively referred to as release factors, RFs). RF1 and RF2 catalyze the first step of translation termination (Bertram et al., 2001), recognizing stop codons and releasing completed peptides from the ribosome (Fig. 1b). RF1 and RF2 directly interact with the ribosome (Capecchi, 1967; Capecchi and Klein, 1970; Petry et al., 2005), and have partially overlapping specificities (Scolnick et al., 1968) (RF1 recognizes stops UAA/UAG, and RF2 stops UAA/UGA). PrmC post-translationally modifies RF1 and RF2, thereby increasing their catalytic activity (Heurgué-Hamard et al., 2002; Mora et al., 2007; Nakahigashi et al., 2002). We targeted translation termination because the resulting downstream changes in expression were anticipated to be modest: given that translation is initiation-limited on most mRNAs (Laursen and Sørensen, 2005), mildly decreasing termination rate was expected to reduce global ribosome availability without altering protein production on a gene-by-gene basis. A better understanding of the physiology of translation termination stress has relevance in synthetic biology, for example in the context of genome-wide stop codon reassignment (Fredens et al., 2019; Johnson et al., 2011, 2012; Lajoie et al., 2013; Wannier et al., 2018).

Through precise measurements of transcriptomes, global translational responses, and cell fitness (defined here as the population growth rate in exponential phase), we elucidate underlying determinants of the RF expression-fitness landscapes (Fig. 1c). We find that idiosyncratic and indirect inductions of regulatory programs are associated with decreases in growth rate in multiple directions of the RF expression subspace. We term such situation ‘regulatory entrenchment,’ whereby the fitness defect caused by perturbing a protein’s expression is strongly and spuriously aggravated by the gene regulatory network. Further, we reconstruct links that connect the initial microscopic perturbation to system-wide effects. At the mechanistic level, we find that occlusion of ribosome binding sites during RF depletion is a common phenomenon leading to changes in expression stoichiometries between adjacent, co-transcribed genes. In particular, we identify the stop codon of a single regulator as a molecular Achilles heel, sensitizing the entire cell to specific RF perturbations. At the regulatory level, we show that removing one gene can mute pleiotropic changes and liberate RF from regulatory entrenchment. Finally, at the systemic level, we report passive proteome compression as a quantitatively tractable cause of growth defects upon massive activation of a regulon.

Our multiscale characterization provides a concrete example of how distal yet focal events triggered by targeted molecular perturbations can have system-wide impacts. Here, the cells’ susceptibility to expression perturbations of specific enzymes is not simply related to the magnitude of the changes in the associated cognate flux, but is instead dictated and amplified by sensitive nodes in the regulatory network. These results underscore the viewpoint that a quantitative understanding of the full system (Boyle et al., 2017; Karr et al., 2012; Liu et al., 2019) might be necessary to predict even qualitatively the shape of expression-fitness landscapes.

## Results

### Linking RF perturbations to changes in genome-wide expression and fitness

We used a combination of genetic tools and high-throughput measurements to map RF expression-fitness landscapes, as well as the underlying mechanistic, regulatory, and systemic responses. We created strains in which two of the three factors (PrmC, RF1, and RF2) can be tuned orthogonally (Fig. S1e-f): PrmC and RF1 in one strain, and PrmC and RF2 in another (with the autoregulatory frameshift removed for RF2 (Craigen and Caskey, 1986; Craigen et al., 1985), Fig. S1a-c). The range of these tunable expression systems spanned 31 to 111-fold and was centered near their respective endogenous levels (Fig. S1d). The global gene expression changes resulting from these perturbations were probed with RNA-seq (Parker et al., 2019) (Methods). Expression levels were converted to fractions of the proteome using ribosome profiling (Ingolia et al., 2009; Li et al., 2014) (Methods for details and assumptions). Ribosome profiling further provided a high-resolution view of translation status in a subset of conditions. In order to precisely measure fitness, defined here as relative population growth rate in exponential phase, we designed a DNA-barcoded competition assay (Parker et al., 2020; Smith et al., 2009) with ±1% precision (±2σ, Fig. S2, Methods). This combination of targeted proteome perturbation with precision transcriptomic and fitness measurements enabled a multiscale assessment of the RF expression-fitness landscapes.

### Perturbing the expression of different RFs decreases fitness through distinct physiological routes

Using our measurement platform, we directly confirmed that endogenous RF expression in exponential phase maximizes growth rate. For conditions at or near endogenous levels of RF1, RF2, and PrmC (dashed lines, Fig. 2e-g), the fitness is indistinguishable from wild-type strains (shaded grey area in Fig. 2e-h corresponds to experimental precision in fitness, |s| < 2σ_s_ = 1.2%, Methods). On the other hand, perturbing RF expression away from endogenous level led to growth defects in most directions of the RF subspace (Fig. 2e-h). Only RF1 overexpression did not cause a measurable decrease in fitness, in part because of the limited maximal achievable level for this particular inducible construct (≈3×, Fig. S1d). These results suggest that RF expression is optimized for exponential phase growth, akin to the expression of several other factors (Dekel and Alon, 2005; Ehrenberg and Kurland, 1984; Li et al., 2014; Parker et al., 2020), and is consistent with the tight evolutionary conservation of expression stoichiometry in biological pathways (Lalanne et al., 2018).

**Fig. 2.**
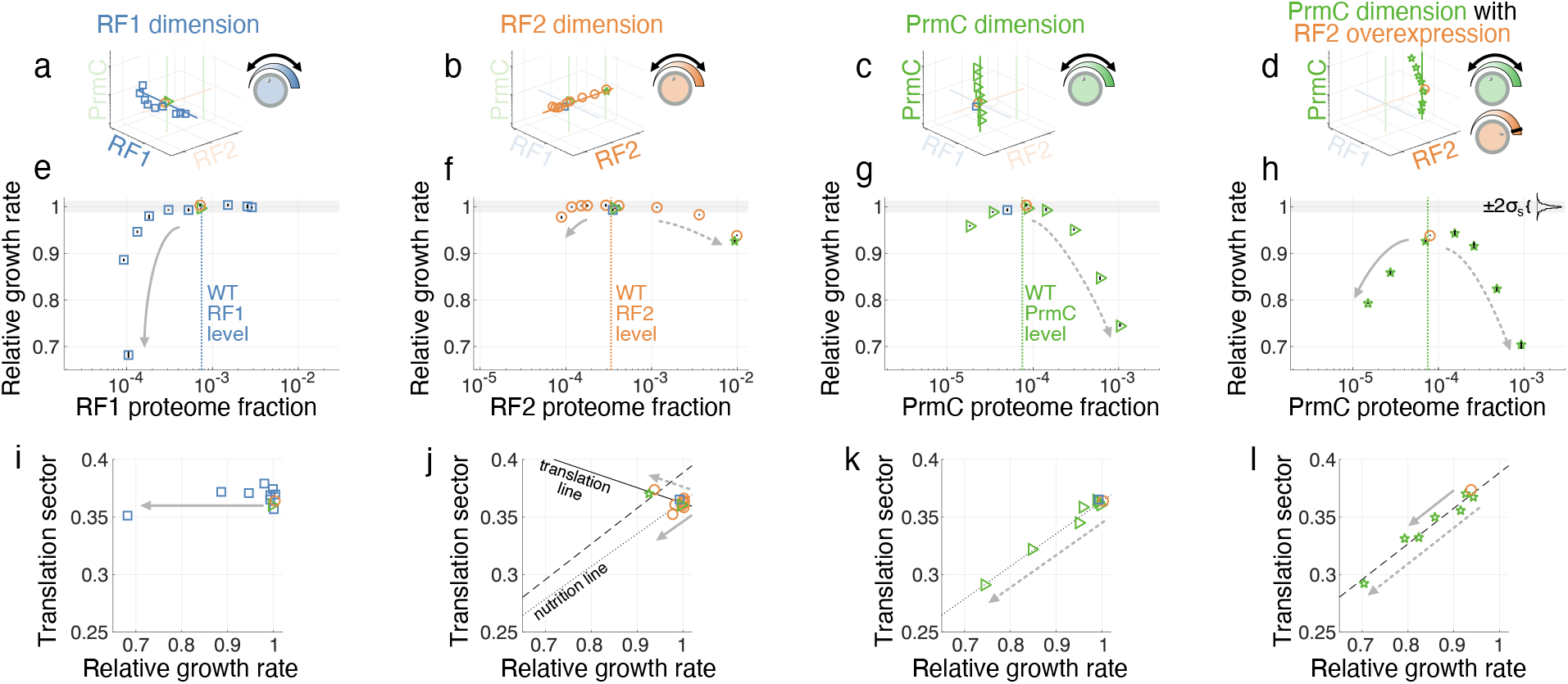
Diverse fitness landscapes and physiological trajectories upon RF expression perturbation. **(a-d)** Profiled orthogonal directions of the RF expression subspace, respectively scanning along the dimension of **(a)** RF1, **(b)** RF2, **(c)** PrmC, and **(d)** PrmC with RF2 overexpression. Axes correspond to expression levels of RF1, RF2, and PrmC. **(e-f)** Cell exponential growth rate measured by competition (relative to wild-type) at corresponding RF levels (reported in units of proteome fraction, derived from a ribosome profiling calibration, Methods) shown in (a-d). Endogenous levels of RFs are indicated with dashed vertical lines. Gray shadings mark the precision of our fitness measurement, defined as ±2σ_s_ = ±1.2%, where σ_s_ is the standard deviation in the measured relative growth rate among isogenic redundantly barcoded strain (see Fig. S2e), with the distribution shown as inset in panel (h). Relative fitness value reported corresponds to the median across isogenic barcoded pairs, and vertical black bars delineate 25^th^ to 75^th^ percentile among such pairs (typically smaller than plot symbol). **(i-l)** Trajectories following RF perturbation in the space of relative growth rate vs. estimated proteome fraction to translation proteins. Matched arrows across panels (e-h) and (i-l) show direction of increasing perturbations in the fitness landscape. Lines in (j-l) correspond to least-square fits. Lines in panels (k) and (l) have been reproduced in panel (j) to highlight of additivity of trajectories under PrmC perturbation with RF2 overexpression. See also Fig. S1-S4.

Away from the optimum, distinct RFs displayed expression-fitness landscapes of different shapes. For example, PrmC has a much more severe growth defect than RF2 at the same levels of overexpression (Fig. 2f vs. 2g), and RF1 knockdown leads to a near-vertical drop in fitness (Fig. 2e), which is more severe than RF2 or PrmC knockdown (though our expression constructs had higher baseline expression for the latter two). Beyond these qualitative differences, our global expression quantification in all these conditions provides a way to assess whether RF-specific fitness defects nevertheless correspond to similar underlying physiological states generic to translation termination defects.

Projecting complex phenotypic data onto low-dimensional manifolds can provide insight on the physiological state underlying growth defects (Hui et al., 2015; Scott et al., 2010, 2014; You et al., 2013). Two cellular-level quantities are of particular interest: (1) the growth rate λ, and (2) the total expression of the mRNA translation machinery, quantified as the summed proteome fraction of translation proteins, termed the translation sector, *ϕ*_R_. Hwa and co-workers have shown (Scott et al., 2010; You et al., 2013; Zhu et al., 2016, 2019) that trajectories in the space of growth λ vs. translation sector *ϕ*_R_ upon a series of increasingly severe perturbations are intimately related to global regulation (Box. 1). In *E. coli*, they found that under numerous ways to inhibit translation, the translation sector *ϕ*_R_ increases concomitantly with a decrease in the growth rate (along the “translation line”, Box 1) as a compensation mechanism (Scott et al., 2010). By contrast, decreasing the nutritional quality of the medium leads to a decrease in growth paralleled by a decrease in the translation sector *ϕ*_R_ (along the “nutrition line”, Box 1), a long-known property in bacterial physiology (Schaechter et al., 1958). Under the bacterial growth laws established in *E. coli* (Box. 1), the states of cells with RF-specific fitness defects were anticipated to collapse on the translation line. In effect, perturbing RF expression was expected to reduce the translation termination rate and thus to reduce ribosome availability.

Surprisingly, the different RF expression perturbations displayed diverse physiological trajectories (Fig. 2i-l). The sharp drop in growth under RF1 knockdown led to little change in the translation sector *ϕ*_R_ (Fig. 2i), analogously to the trajectories associated with non-translation targeting antibiotics (Scott et al., 2010). By contrast, the growth defects upon PrmC expression perturbations were associated with a decrease in the translation sector (Fig. 2k-l), leading to movement along the nutrition line, which was unexpected given the fixed quality of the growth medium. RF2-perturbed cells moved slightly in different directions following overexpression (translation line) and knockdown (nutrition line, although high basal activity of our expression construct limited the magnitude accessible growth defects) (Fig. 2j). Interestingly, physiological changes were additive under combined RF perturbations: tuning PrmC expression in combination with RF2 overexpression led to movement along the nutrition line, but shifted by the movement along the translation line by the RF2 perturbation (dashed vs. dotted lines in Fig. 2j-l, Supplementary Discussion).

The physiological divergences observed for the different RFs perturbations were intriguing given the similar involvement of these three proteins in translation termination, and pointed to possible RF-specific pleiotropic effects. Analyzing the transcriptomes following perturbations revealed distinct responses along the orthogonal RF expression directions. RF1 knockdown led to changes in motility and biofilm genes (Fig. S4c-d), PrmC perturbations massively induced the general stress σ^B^ regulon (Fig. 3a-b, S3a-d), and RF2 perturbation led to little changes across the full range of expression, except for a modest σ^B^ induction at maximal knockdown (Fig. S3o-p). Together, these results suggest that each RF has a mechanistically distinct relationship between expression and fitness.

**Fig. 3.**
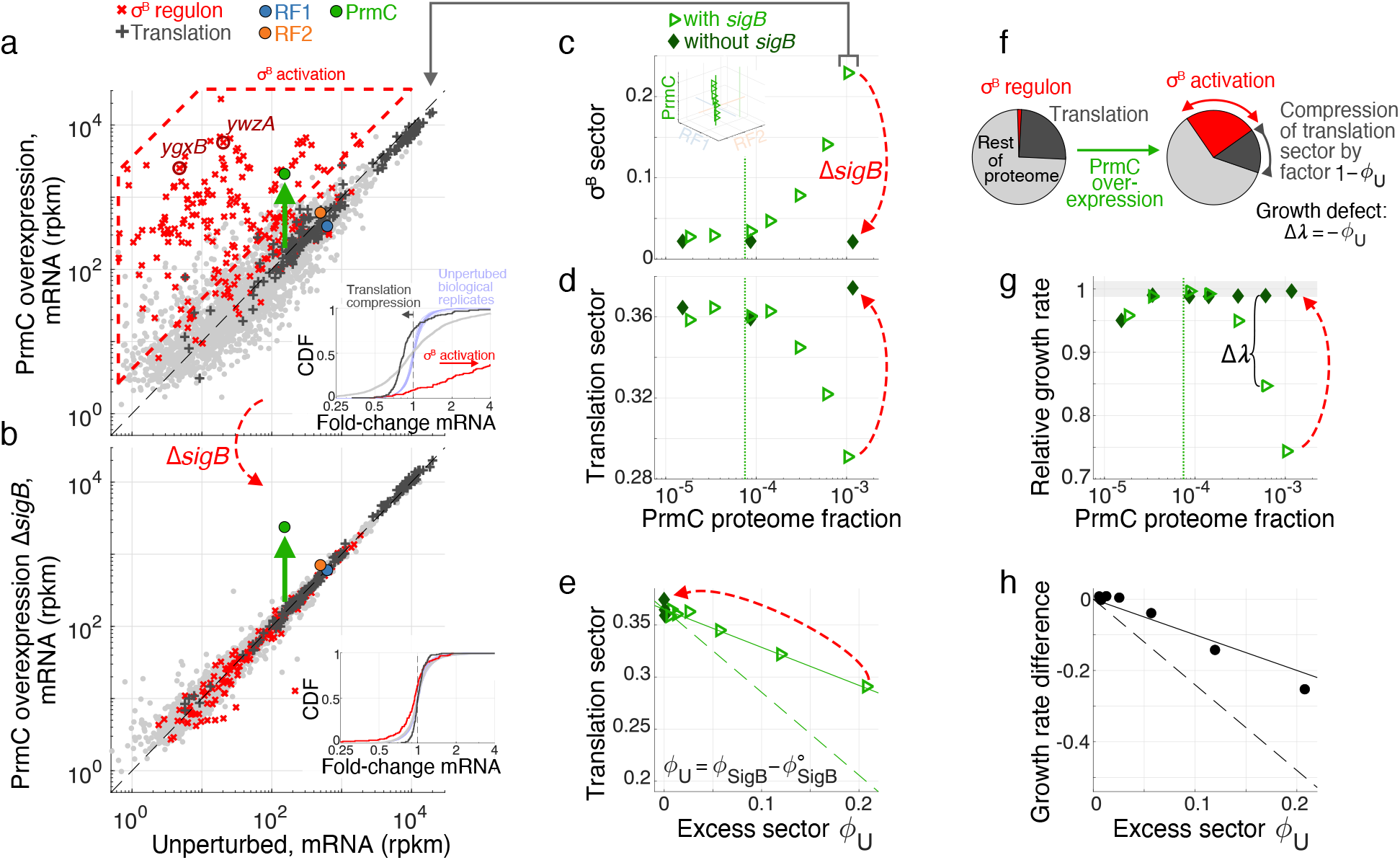
σ^B^ regulon induction upon PrmC overexpression compresses the translation sector. **(a)** mRNA levels (reads per million mapped reads per kilo-base, rpkm, genes with >5 reads mapped shown) for maximal PrmC overexpression (green arrow) versus unperturbed cells (average across control datasets, Methods). σ^B^ regulon members and translation related proteins are marked in red **×** and dark grey **+** respectively. Targets for RT-qPCR measurements of σ^B^ induction (*ygxB*, *ywzA*, Fig. 5) are highlighted in dark red. mRNA levels for RF1 (blue), RF2 (orange), and PrmC (green), are marked by dots (corrected for translation efficiency of ectopic expression constructs, Methods). σ^B^ regulon activation is marked by dashed red polygon. Inset shows cumulative distribution of fold-changes in mRNA levels (red σ^B^ regulon, dark gray translation, pale gray rest of proteins). σ^B^ activation and translation compression are highlighted by arrows indicating shift in median expression. As a comparison, distribution of fold-changes for all genes among unperturbed replicates are show in light blue. **(b)** Same as (a), but with a deletion of gene *sigB*, which abrogates σ^B^ regulon activation, and restores genome-wide expression levels despite PrmC overexpression (light gray line in inset). **(c)** Quantification of the proteome fraction to the σ^B^ regulon (σ^B^ sector) as a function of PrmC (see Methods for calibration from transcriptome to proteome fraction). Dashed vertical line marks endogenous PrmC level. Inset reproduces broader context of data in RF expression subspace (Fig. 2c). In panels (c-e) and (g), open light green triangles correspond to cells with *sigB*, and filled dark green diamonds to cells without *sigB* (deletion). **(d)** Similar to (c), but quantifying proteome fraction to the translation sector. **(e)** Proteome fraction of the translation sector as a function of the excess proteome fraction to the σ^B^ regulon, denoted *ϕ*_*U*_. Dashed line corresponds to growth laws prediction (Box. 1, Eq. 1, using parameters *κ*_*n*_, *κ*_*t*_, *ϕ*_∘_, and 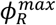 obtained from fits in Fig. 2j-k), full line corresponds to decrease by factor 1 − *ϕ*_*U*_. **(f)** Schematic illustration of passive proteome fraction compression under σ^B^ activation (increase in regulon expression). **(g)** Relative growth rate as a function of PrmC level, with and without *sigB*. **(h)** Growth difference with and without *sigB* (corresponding to Δλ panel (g)), as a function of excess proteome fraction to the σ^B^ regulon. Dashed lines are growth laws prediction (Box 1, Eq. 2), full line corresponds to −*ϕ*_*U*_. See also Fig. S3 and S4.

### Fitness defects during PrmC overexpression can be traced back to regulatory response by σ^B^ activation

To further identify the origin of some of the observed growth defects, we focused on perturbations to PrmC. The growth defect following PrmC overexpression (Fig. 2g) coincided with dramatic activation of the general stress regulon, which is driven by the alternative sigma factor σ^B^ (gene *sigB*) (Hecker et al., 2007; Helmann et al., 2001; Petersohn et al., 2001; Price et al., 2001; Price CW, 2002; Zhu and Stülke, 2018), from 2% basal proteome synthesis fraction in wild-type to >20% (Fig. 3a, c, see Methods for ribosome profiling calibration). This increase in the expression of σ^B^ genes was accompanied by a corresponding decrease in the translation sector *ϕ*_R_ (Fig. 3d-e), which raised the possibility that a substantial portion of the growth defect following PrmC overexpression was associated with the physiological burden of expressing >100 σ^B^-dependent genes at high levels (Benson and Haldenwang, 1992; Bernhardt et al., 1997; Boylan et al., 1992). Alternatively, induction of the stress regulon might have directly responded to, and played a role in mitigating, translational stress resulting from PrmC expression changes. As additional evidence of systemic stress, PrmC overexpression leads to global changes in gene expression beyond the σ^B^ regulon (broadening of fold-change distribution, Fig. 3a inset). Whether these global changes were the result of the PrmC perturbation, or secondary to σ^B^ activation, was to be clarified.

To determine whether σ^B^ exacerbates or mitigates fitness defects under PrmC perturbations, we profiled RF-inducible cells lacking the *sigB* gene (Δ*sigB*). As expected, deleting *sigB* completely abrogated regulon induction (Fig. 3b-c, dashed red arrows), but surprisingly also restored genome-wide expression to near endogenous levels (Fig. 3b inset). Hence, global expression changes (gray line in inset Fig. 3a) upon PrmC overexpression were caused by σ^B^ activation and its downstream consequences (e.g., sigma factors competition (Farewell et al., 1998)). Importantly, the PrmC overexpression growth defect was completely rescued following *sigB* deletion (Fig. 3g dashed red arrow), concomitantly with the restoration of the translation sector to endogenous levels (Fig. 3d-e dashed red arrows). These observations were reproducible under simultaneous PrmC/RF2 overexpression (Fig. S3a-i, Supplementary Discussion).

Taken together, these results suggest that the gene *sigB* constitutes a *trans*-modifier (Hou et al., 2019) of the PrmC expression-fitness landscape. Through its activation, σ^B^ drastically exacerbates the cell’s sensitivity to expression perturbation of distal gene *prmC*. Overall, the induction of the general stress regulon has no measured benefit in the context of PrmC perturbations. Gene *sigB* therefore ‘entrenches’ PrmC expression by imposing a massive physiological burden to cells that is in addition to, and much larger than, the effect directly related to perturbing PrmC.

### Systemic proteome compression by σ^B^ explains fitness defect

We hypothesized that the growth defect resulting from σ^B^ activation mainly arose from the cost of expression of regulon genes. The burden of expressing proteins has been extensively characterized in *E. coli* (Andrews and Hegeman, 1976; Dong et al., 1995; Scott et al., 2010; Stoebel et al., 2008). Gratuitous expression of genes causes “proteome compression”, whereby the abundances of other proteins globally decrease in response to the finite total synthesis capacity of the cell (Scott et al., 2010). The resulting reduction in the translation sector, responsible for generating biomass, leads to a growth defect, which would be in addition to any protein-specific effects such as misfolding or aggregation. Bacterial growth laws, established in *E. coli*, predict the quantitative relationship between proteome fraction of gratuitous proteins and growth rate under this proteome compression model (Box 1).

Despite the complexity of the induced regulon (>100 diverse genes) and the resulting secondary changes in gene expression (Fig. 3a), σ^B^ activation did lead to simple compression of the translation machinery (Fig. 3e-f). Indeed, the translation sector (full line, Fig. 3e, S3g) and growth rate (full line, Fig. 3h, S3i) linearly decreased in a one-to-one proportion with the increase in excess proteome fraction to the σ^B^ regulon. Interestingly, we observed a smaller growth defect than expected from the *E. coli* growth law (dashed vs. full lines, Fig. 3e, 3h, S3g, S3i, Box 1). Further assessment of the generality of growth laws to species other than *E. coli* will constitute promising future research avenues.

Compression of the translation sector due to σ^B^ activation provides a systemic explanation for the movement along the nutritional line under PrmC perturbation (Fig. 2k-l). Abrogation of this effect following *sigB* deletion (Fig. 3d-e, S3f-g, dashed arrow) highlights that physiological trajectories can be modulated by individual regulatory proteins, as has been also demonstrated for tRNA synthetases and the stringent response in *B. subtilis* (Parker et al., 2020). However, unlike the stringent response, which is cognate to synthetase perturbation and beneficial to cell growth, the σ^B^ response here is detrimental.

### Activation of regulator σ^B^ does not result from general stress-sensing mechanisms

The detrimental effect on fitness of σ^B^ activation under RF stress (Fig. 3g) suggests that its induction is idiosyncratic in this context, as opposed to being the result of a cognate sensing mechanism. We hypothesized that RF-specific perturbations led to changes in of a small number of proteins that ultimately activated σ^B^. In support of this view, σ^B^ is only activated when certain RFs are perturbed: varying RF1 expression over the full achievable range (0.1× to 3× endogenous) did not change σ^B^ regulon expression (Fig. S4c), whereas RF2 knockdown, but not overexpression, was associated with σ^B^ activation (Fig. S3o-p, and lowering basal RF2 expression further increased σ^B^ activation, see Supplementary Data 8). Similar to PrmC overexpression, the fitness defects under RF2 knockdown was rescued by *sigB* deletion (Fig. S3q), also arguing for regulatory entrenchment of RF2 expression by σ^B^.

To confirm that σ^B^ activation arose from perturbations specific to distinct RFs and not from general translation stress-sensing mechanisms, we used ribosome profiling to quantify translation in cells acutely depleted of RFs. We used CRISPRi transcriptional interference (Peters et al., 2016; Qi et al., 2013) to separately knockdown RF1/PrmC (co-transcribed) and RF2. In both cases, we found evidence of translational stress in the forms of idle ribosomes at the corresponding stop codons and queuing upstream (Fig. 4a), similar to what was previously observed under different translation termination stresses (Baggett et al., 2017; Mangano et al., 2020; Saito et al., 2020). Specifically, we observed queues at UAG stops under RF1/PrmC depletion, queues at UGA stops under RF2 depletion, and no queues in wild-type, or at UAA stops in either depletion. Queues were longer on mRNAs with high translation efficiency, as expected from models of stochastic queuing (Baggett et al., 2017; Bergmann and Lodish, 1979; Mitarai et al., 2008; Saito et al., 2020) (Fig. S5b-c, S5h, Supplementary Discussion). The severity of the translation stress, as assessed by the magnitude of accumulation of ribosomes on stop codons, was similar for the PrmC/RF1 and RF2 depletions (Fig. 4a). Importantly however, a robust σ^B^ activation was only observed under RF2, and not RF1/PrmC, knockdown (Fig. 4b-c). The simultaneous presence of ribosome queues (Fig. 4a, middle row) and absence of σ^B^ activation under RF1/PrmC depletion (Fig. 4b) confirmed that σ^B^ induction was not channeled through a sensor of translation stress agnostic to the identity of ribosome-jammed mRNAs. σ^B^ induction was instead likely due to gene-specific expression changes driven by RF stop codon specificities.

**Fig. 4.**
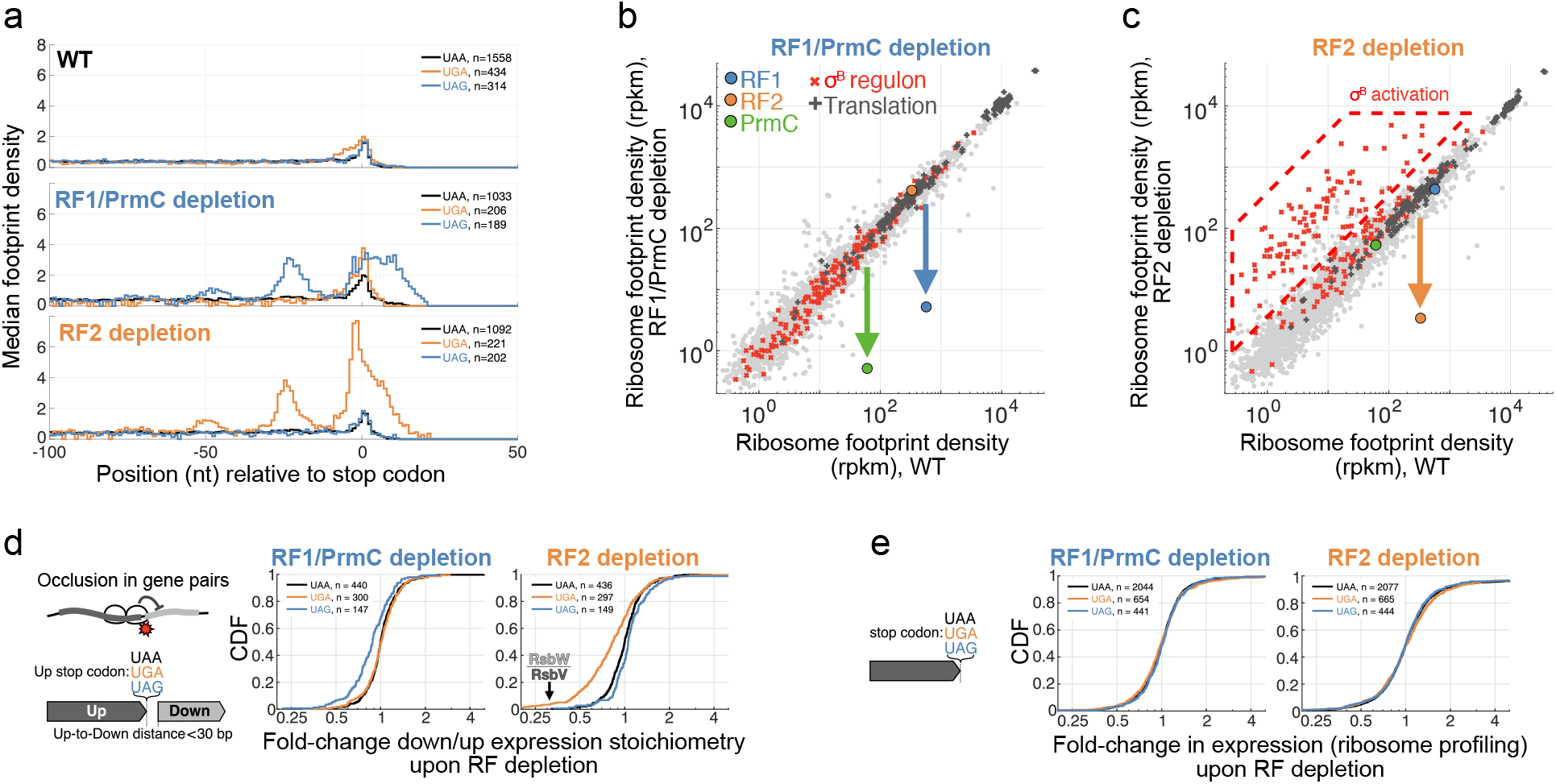
Translation termination stress and gene expression changes upon acute RF depletion with CRISPRi. **(a)** Metagene trace (Methods) of gene-normalized ribosome footprint read density (center-mapped) upstream of stop codons stratified by stop codon. Wild-type (top), RF1/PrmC depletion (middle), and RF2 depletion (bottom) are shown. Ribosome queues upstream of stop codon cognate to RF perturbations can be seen. **(b-c)** Comparison of expression (ribosome footprint density, rpkm) for **(b)** RF1/PrmC depletion, and **(c)** RF2 depletion, vs. wild-type. Regulons and RF are marked as in Fig. 3a. RF2 depletion leads to σ^B^ activation, in contrast to RF1/PrmC depletion. **(d)** Fold-change in the downstream-to-upstream expression (ribosome profiling) stoichiometry upon RF depletion for co-directional genes within 30 bp, and stratified by the stop codon of the upstream gene. A mild but systematic and stop-codon specific decrease in downstream gene expression is seen (RF1/PrmC: median UAG fold-change compared to UAA (FC_UAG_)= 0.88, p<10^−6^; RF2: FC_UGA_ = 0.82, p<10^−6^, p-values from stop codon reshufflings, Methods). We ascribe this effect to obstruction of downstream ribosome initiation by idle upstream terminating ribosomes. Some proteins pairs, such as RsbW/RsbV, are affected strongly. **(e)** Fold-change in expression between RF depletion and wild-type stratified by stop codon of genes. No systematic effect for the different stop codons is seen (RF1/PrmC: FC_UAG_ = 1.01, p=0.69; RF2: FC_UGA_=1.02, p=0.86), indicating lack of strong change in expression for genes with compromised translation termination. See also Fig. S3, Fig. S5.

### RF depletion mechanistically leads to stoichiometric imbalance of co-transcribed genes

We hypothesized that a consequence of RF depletion is that the ribosomes idling at the corresponding stop codons would sterically occlude translation initiation of downstream genes. Because bacterial genes are often closely spaced in polycistronic operons, elevated ribosome occupancy at stop codons may prevent other ribosomes from binding to the next gene (Fig. 4d). Consistent with this hypothesis, we found that among co-directional genes separated by a ribosome footprint or less (≤30 nt), the downstream genes exhibit a mild but stop-specific decrease in expression when the cognate RF is depleted (Fig. 4d, RF1/PrmC: FC_UAG_ = 0.88, p<10^−6^; RF2: FC_UGA_ = 0.82, p<10^−6^, see Fig. S5d-f for control groups). This result differs from a lack of effects reported in *E. coli* during knockdown of ribosome recycling factors (RF4), which act on ribosomes post RF1/RF2-mediated peptide release. These ribosomes may have different mobility and re-initiation properties (Saito et al., 2020). Although the effect size was modest overall, some gene pairs were affected more heavily. For example, highly translated upstream genes are correlated with stronger decrease in downstream expression Fig. S5i. Gene pairs joined by <u>A**UGA**</u>, i.e., the most common overlapping start and stop punctuation (Fig. S5j) possibly involved in translational coupling, did not show a substantial difference (Fig. S5k). Overall, these genome-wide measurements point to occlusion of ribosome binding sites by upstream termination-idle ribosomes as a cause of RF-specific gene expression changes.

In contrast to the effect on co-transcribed genes downstream, the expression of genes ending with the compromised stop codons were not substantially affected either at the level of translation (Fig. 4e, RF1/PrmC: median UAG fold-change compared to UAA (FC_UAG_)= 1.01, p=0.69; RF2: FC_UGA_=1.02, p=0.86; p-values from stop codon reshufflings, Methods), or mRNA (Fig. S5g, RF1/PrmC: FC_UAG_=0.98, p=0.06; RF2: FC_UGA_=0.98, p=0.02). This lack of systematic changes in expression for genes with translation termination defects is consistent with the observed ribosome queues being too short to interfere with translation initiation at the upstream start of the ORF (queue size of <4 stalled ribosomes globally, Fig. 4a) (Bergmann and Lodish, 1979; Jin et al., 2002). This result also suggests that translational quality control and surveillance mechanisms, such as mRNA cleavage at empty A-site (Hayes and Sauer, 2003; Li et al., 2007) and ribosome collision sensors (Ferrin and Subramaniam, 2017), play a minor role in comparison to start codon occlusion for downstream genes in the current context.

### Mechanism for σ^B^ activation by hypersensitivity of a single cis-element

Expression stoichiometry of genes *rsbW*/*rsbV*, whose ORFs overlap via the RF2 stop codon (−4 nt overlap <u>A**UGA**</u>, Fig. 5b, Fig. S5j), was one of the most sensitive to RF2 knockdown (3.0× change, Fig. 4d, compared to 1.1× in RF1/PrmC knockdown). Intriguingly, these two proteins form the regulatory core of σ^B^ activation. Upstream signaling events converge on a phosphorylation-dependent partner-switching mechanism involving the anti-sigma factor RsbW and the anti-anti-sigma factor RsbV, which respectively inhibits and activates σ^B^ (Pané-farré et al., 2017). As additional control points, RsbW can deactivate RsbV by direct phosphorylation (Hecker et al., 2007; Price CW, 2002) and the three genes *rsbW*, *rsbV*, and *sigB* are co-transcribed under a σ^B^ dependent promoter (among other transcript isoforms) (Fig. 5a). These interlocked positive and negative feedbacks (Fig. 5a) render σ^B^ activation hypersensitive to the stoichiometry of its regulators RsbV and RsbW (Igoshin et al., 2007; Locke et al., 2011).

**Fig. 5.**
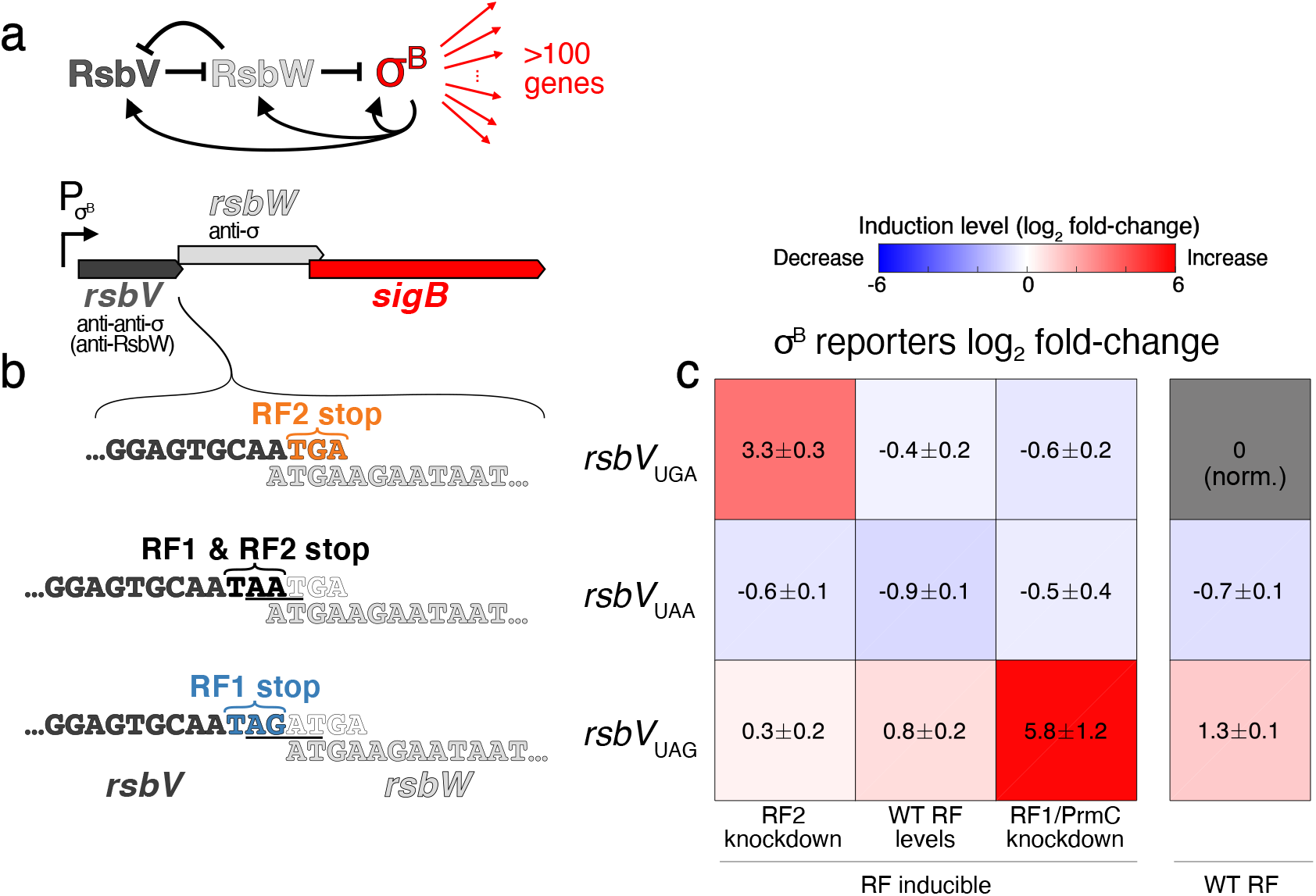
Swapping a single stop codon rewires σ^B^ induction susceptibility to RF perturbation. **(a)** Schematic of simplified σ^B^ regulatory architecture and operon structure. **(b)** Different *rsbV* stop codon variants considered. Top row shows endogenous configuration *rsbV*_UGA_, with RF2 dependent stop codon and <u>A**UGA**</u> open reading frames overlap. Middle row shows the UAA RF agnostic variant *rsbV*_UAA_, obtained by adding AAT (underlined). Bottom row shows the RF1 dependent UAG allele *rsbV*_UAG_, obtained by inserting AGAT (underlined). **(c)** Average of reporter gene (2-3 biological replicates for each allele/condition, ± indicates standard deviation of the mean over replicates) log_2_ fold-change compared to wild-type for σ^B^ reporter genes (*ywzA* and *ygxB*, highlighted in Fig. 3a) as quantified by RT-qPCR (raw data in Supplementary Data 8, Methods) showing strong induction of reporter genes in the RF perturbations cognate to the stop codon of the *rsbV* variant. Rows correspond to *rsbV* stop variant in panel (b), and columns to RF expression conditions. RF inducible measurements were done in strains with orthogonally tunable RF1/PrmC (IPTG), and RF2 (xylose) (Methods). See also Fig. S6.

Given that we observed σ^B^ activation in RF2, but not RF1/PrmC, knockdown (Fig. 4c vs 4b), and that the expression of σ^B^ regulators RsbV/RsbW was affected in these conditions (Fig. 4d), we hypothesized that termination defects at the *rsbV* UGA (RF2-specific) stop codon was the cause of the σ^B^ activation following RF perturbations. As such, switching the *rsbV* stop codon was predicted to be sufficient to rewire the susceptibility of σ^B^ activation to different RF perturbation. For example, with a switched UAG stop for gene *rsbV*, σ^B^ might be activated upon RF1, but not RF2, knockdown.

To test whether σ^B^ activation sensitivity was indeed focally dependent on this *cis*-element, we generated a set of *rsbV* stop codon variants (UAA: RF1 and RF2 activity, UAG: RF1 only activity) (Fig. 5b). These variants were cleanly inserted at the endogenous *sigB* operon in wild-type and in an RF inducible strain (RF1/PrmC and RF2, with frameshift removed, under orthogonal promoters, Methods). Under various RF expression conditions (endogenous, RF2 knockdown, RF1/PrmC knockdown), we monitored mRNA levels of two responsive σ^B^-dependent genes by RT-qPCR (*ywzA* and *ygxB*, highlighted in Fig. 3a, Methods). Basal σ^B^ activity at endogenous RF expression differed slightly for the different *rsbV* stop variants, likely as a result of perturbed ORF positioning. Strikingly, we observed strong σ^B^ activation only in strains with the *rsbV* stop cognate to the RF perturbation (Fig. 5c, raw data in Supplementary Data 8). Consistently with previous experiments, RF2 knockdown led to an increase (>8×) in the expression of our σ^B^ reporter genes in the *rsbV*_UGA_ variant (Fig. 5c, top row). In addition, induction of reporter genes was observed for the *rsbV*_UAG_ variant only under RF1/PrmC knockdown (>50×, Fig. 5c, bottom row). No induction in either RF perturbation was observed for the *rsbV*_UAA_ variant (Fig. 5c, middle row), in accordance with the UAA stop codon being RF agnostic. The *rsbV*_UAA_ variant was confirmed functional (>500× induction, indistinguishable from the wild-type *rsbV*_UGA_ variant following 5 minutes of 4% v/v ethanol stress, Supplementary Data 8). Hence, our stop codon swapping experiment demonstrates that systems-wide susceptibility to a distal molecular perturbation can be encoded through a single sensitive *cis*-element, and that such susceptibility can be entirely rewired with minimal genetic changes.

## Discussion

We have shown that quantitative genome-wide measurements in conjunction with precision fitness quantification can be used to reconstruct chains of events across biological scales relating a molecular perturbation (change in expression of a specific protein) to a whole cell phenotype (growth rate). We find that in *B. subtilis*, important portions of the selective pressures on peptide release factor abundances is through idiosyncratic activation of distal endogenous regulatory programs (Fig. 6). By analogy with interactions between residues constraining the evolution of individual proteins (Bridgham et al., 2009; Pollock et al., 2012; Shah et al., 2015; Starr et al., 2018), we term this strong dependence of enzymes’ expression-fitness landscapes on the network of genetic interactions ‘regulatory entrenchment’ (Fig. 6).

**Fig. 6.**
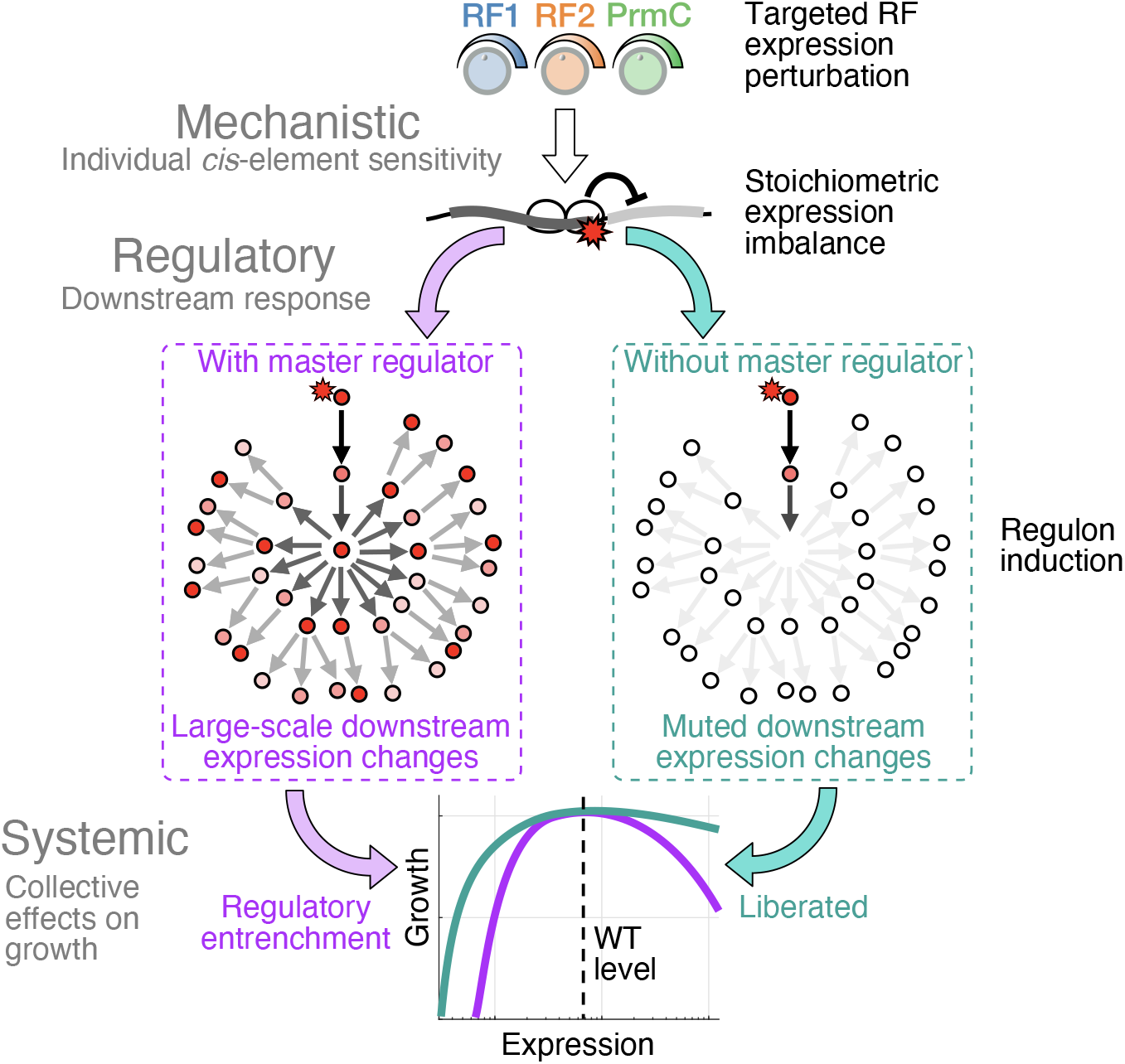
Regulatory entrenchment in gene expression-fitness landscapes. Perturbation of specific enzyme expression can lead to changes in the expression of a subset of genes through the sensitivity of *cis*-regulatory elements. These expression imbalances can further reverberate in the network of regulatory interactions in the cell through focal points such as master regulators, which control the expression of multiple genes (purple box). In the absence of these regulators, downstream responses are muted (teal box). Pleiotropic expression changes exacerbate the fitness defect of perturbing the original enzyme. The cell’s susceptibility to expression is then not simple relation of the decreased enzymatic flux, but also depends on the aggregated impacts regulatory susceptibilities of the whole cell. Cells can be liberated from regulatory entrenchment by removing nodes in the network. See also Fig. S3 and S4.

Despite our focus on translation termination factors, which we anticipated to have limited impacts on gene expression (translation termination being non-limiting on mRNAs), pleiotropic effects related to activation of specific regulons still dominated the connection between RF expression and fitness. As an example of molecular determinants shaping expression-fitness landscapes, we found that a single *cis*-element (identity of the *rsbV* stop codon, Fig. 5b-c) mechanistically led to RF-specific expression imbalance (altered stoichiometry of σ^B^ regulators RsbW and RsbV, Fig. 4d). This imbalance was amplified to global transcriptome remodeling through to the activation of σ^B^ (Fig. 3a, S3c). These changes ultimately had systemic impacts (growth defects by proteome compression, Fig. 3e, S3g, S3o). Although the exact cause of σ^B^ activation due to PrmC overexpression remains to be determined, deletion of the *sigB* gene was sufficient to flatten the landscape in two directions of the RF expression space (PrmC overexpression Fig. 3g, S3h and RF2 knockdown Fig. S3q).

The adaptive nature of σ^B^ activation upon RF-stress remains a possibility, but the lack of corresponding fitness advantage in our experiments suggests otherwise (Fig. 3g, S3h). Bioinformatic analysis of homologous σ^B^ operons shows some conservation in the *rsbV/rsbW* ORF overlap (Fig. S6, Methods). Overlap of these genes could however be driven by the requirement of a precisely tuned expression stoichiometry (Igoshin et al., 2007; Locke et al., 2011), enabled by translational coupling via the −4 nt start/stop overlap configuration <u>A**UGA**</u>. In such scenario, susceptibility to RF perturbation would emerge as an evolutionary byproduct of selective pressures that operate on other features of the system (Gould, 1978).

As an additional example of regulatory entrenchment in our data, the sharp decrease in fitness upon RF1 knockdown (Fig. 2e) was associated with gene expression changes characteristic of the motile-to-sessile bistable switch in *B. subtilis* (Chai et al., 2009; Kearns and Losick, 2005) (Fig. S4). Given the complexity of the regulation of motility genes (Mukherjee and Kearns, 2014), we have not been able to identify a unique plausible molecular link to RF1 knockdown (Supplementary Discussion). The relationship between the transition to sessile state and the plummeting cell growth rate also remains elusive. These underscore the challenges of reconstructing causal features underlying expression-fitness landscapes, even with detailed under-the-hood information about the cell’s internal state. Our case study highlights that exhaustive information at multiple levels (Fig. 1a, inset ii), down to the response of all *cis*-regulatory elements, might be required to achieve fully predictive models given how easily perturbations propagate across scales in biological systems. A lack of global characterization of the susceptibilities and interaction of components in regulatory networks currently constitute an important bottleneck in systems biology, although at-scale methods are being developed to dissect genetic networks (Calderon et al., 2020; Muller et al., 2020).

We anticipate that master regulators will be common focal points of regulatory entrenchment, as seen here with general stress factor σ^B^ and motility factor σ^D^, due to the responsiveness of their activation and their direct impact on a large number of genes. As such, successful predictions will presumably be more easily achieved in near-minimal organisms with limited regulation (Baby et al., 2018; Hutchison et al., 2016). Given the broadly conserved expression stoichiometry of enzymes across evolution (Lalanne et al., 2018), we speculate that endogenous expression levels satisfy generic optimization principles, such as flux maximization under resource allocation constraints (Kurland and Ehrenberg, 1987). However, since diverse bacteria harbor distinct *cis*-elements, operonic organization, and regulatory architectures, the shape of expression-fitness landscapes away from the optima might generally differ as a result of regulatory entrenchment.

## Supporting information

Methods with supplementary figures

## Acknowledgments

We thank J. M. Peters for providing CRISPRi strains, members of the G.-W.L. and A. Grossman labs for critical discussions, and T. Starr for introducing us to the concept of entrenchment. We thank L. Herzel for critical comments on the manuscript. This research was supported by NIH grant R35GM124732, the NSF CAREER Award, the Smith Odyssey Award, the Pew Biomedical Scholars Program, a Sloan Research Fellowship, the Searle Scholars Program, the Smith Family Award for Excellence in Biomedical Research, NSERC doctoral Fellowship and HHMI International Student Research Fellowship to J.-B.L., NIH T32GM007287 to D.P.

## Author contributions

J.-B.L. and G.-W. L. designed experiments and analysis; J.-B.L. performed experiments, collected data and performed analysis; J.-B.L. and D.P. designed and optimized competition experiments; D.P. established defined growth medium for *B. subtilis* and RNA-seq method; J.-B.L. and G.-W. L. wrote the manuscript with comments from D.P.

## Box 1. Bacterial growth laws

The bacterial growth laws (Scott et al., 2010, 2014; You et al., 2013) constitutes an empirical framework recapitulating the global regulatory architecture of *E. coli* under nutrition and translation stress in steady-state growth. The growth laws connect the growth rate λ to the abundance of coarse-grained proteome sectors (translation or “ribosome affiliated” proteins *ϕ*_*R*_; unregulated proteins which includes metabolic enzymes *ϕ*_*P*_, and gratuitous unnecessary non-toxic proteins *ϕ*_*U*_) through three mathematical relationships:

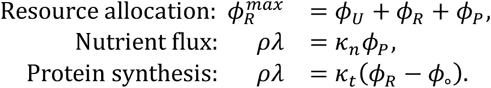

Above, phenomenological parameters *ϕ*_∘_ corresponds to an inactive ribosome fraction, *κ*_*n*_ is the nutritional capacity, *κ*_*t*_ is the translational capacity (proportional to the rate of protein synthesis), and *ρ* is a fixed conversion factor. Nutritional and translational capacities are defined through slopes of the lines along physiological trajectories (Fig. Box. 1).

The growth laws can be interpreted as amino acid flux conservation in steady-state growth: the flux generated by catabolic enzymes and transporters (*ϕ*_*P*_) matches the biomass production flux by translation proteins (*ϕ*_*R*_).

**Fig. Box 1:**
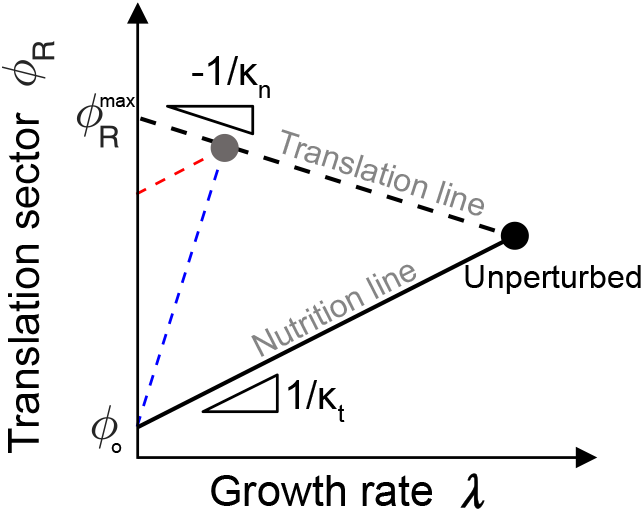
Schematic summary bacterial growth laws. Translation sector (proteome fraction to translation proteins) vs. growth rate. Upon change in growth medium quality or overexpression of unnecessary non-toxic proteins, the cell’s physiological state moves along 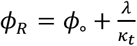 (nutrition line). Following perturbation to translation, trajectories follow 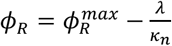 (translation line). Composition of nutritional and translation perturbation can in principle be used to identify nature of perturbation to translation (increase in inactive ribosome, red line vs. decrease in translational capacity, blue line).

Changing the medium quality changes nutritional capacity *κ*_*n*_ without altering the translational *κ*_*t*_ and inactive ribosome content *ϕ*_∘_. Then, the translation sector *ϕ*_*R*_ decreases following the nutrition line:

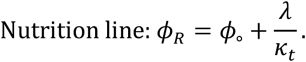

In contrast, in a fixed medium (fixed *κ*_*n*_), but upon perturbation to translation (either through the increasing the fraction of inactive ribosome *ϕ*_∘_, e.g., by chloramphenicol treatment (Dai et al., 2016; Scott et al., 2010), or decreasing translation rate *κ*_*t*_, e.g., by fusidic acid treatment (Zhu et al., 2019)), the translation sector increases following:

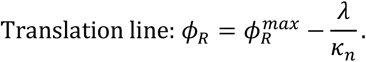

Upon expression of unnecessary proteins to proteome fraction *ϕ*_*U*_, the growth laws above predict movement along the nutrition line (since *κ*_*t*_ and *ϕ*_∘_ are assumed fixed), and a fractional decrease in the translation sector and growth:

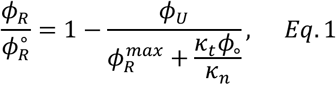

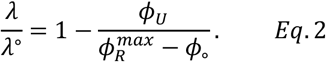

The slope of *ϕ*_*R*_ vs. *ϕ*_*U*_is determined by, which relates to the posited incompressibility of fraction 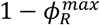 of the proteome. The different slopes (Fig. 3e, 3h, S3g, S3i) we observe in *B. subtilis* suggest possible differences in the growth laws in this species.

## References

Agarwal, V., and Shendure, J. (2018). Predicting mRNA abundance directly from genomic sequence using deep convolutional neural networks. BioRxiv 31, 416685.

Andrews, K.J., and Hegeman, G.D. (1976). Selective disadvantage of non-functional protein synthesis in Escherichia coli. J. Mol. Evol. 8, 317–328.

Baby, V., Lachance, J.-C., Gagnon, J., Lucier, J.-F., Matteau, D., Knight, T., and Rodrigue, S. (2018). Inferring the Minimal Genome of Mesoplasma florum by Comparative Genomics and Transposon Mutagenesis. MSystems 3, 1–14.

Baggett, N.E., Zhang, Y., and Gross, C.A. (2017). Global analysis of translation termination in E. coli. PLoS Genet. 1–27.

Benson, A.K., and Haldenwang, W.G. (1992). Characterization of a regulatory network that controls σ(B) expression in Bacillus subtilis. J. Bacteriol. 174, 749–757.

Bergmann, J.E., and Lodish, H.F. (1979). A Kinetic Model of Protein Synthesis. 254, 11927–11937.

Bernhardt, J., Völker, U., Völker, A., Antelmann, H., Schmid, R., Mach, H., and Hecker, M. (1997). Specific and general stress proteins in Bacillus subtilis - A two-dimensional protein electrophoresis study. Microbiology 143, 999–1017.

Bertram, G., Innes, S., Minella, O., Richardson, J.P., and Stansfield, I. (2001). Endless possibilities: Translation termination and stop codon recognition. Microbiology 147, 255–269.

de Boer, C.G., Vaishnav, E.D., Sadeh, R., Abeyta, E.L., Friedman, N., and Regev, A. (2020). Deciphering eukaryotic gene-regulatory logic with 100 million random promoters. Nat. Biotechnol. 38, 56–65.

Bogard, N., Linder, J., Rosenberg, A.B., and Seelig, G. (2019). A Deep Neural Network for Predicting and Engineering Alternative Polyadenylation. Cell 178, 91–106.e23.

Boylan, S.A., Rutherford, A., Thomas, S.M., and Price, C.W. (1992). Activation of Bacillus subtilis transcription factor σ(B) by a regulatory pathway responsive to stationary-phase signals. J. Bacteriol. 174, 3695–3706.

Boyle, E.A., Li, Y.I., and Pritchard, J.K. (2017). An Expanded View of Complex Traits: From Polygenic to Omnigenic. Cell 169, 1177–1186.

Bridgham, J.T., Ortlund, E.A., and Thornton, J.W. (2009). An epistatic ratchet constrains the direction of glucocorticoid receptor evolution. Nature 461, 515–519.

Calderon, D., Ellis, A., Daza, R.M., Martin, B., Tome, J.M., Chen, W., Chardon, F.M., Leith, A., Lee, C., Trapnell, C., et al. (2020). TransMPRA: A framework for assaying the role of many trans-acting factors at many enhancers 2 3. BioRxiv 2020.09.30.321323.

Cambray, G., Guimaraes, J.C., and Arkin, A.P. (2018). Evaluation of 244,000 synthetic sequences reveals design principles to optimize translation in escherichia coli. Nat. Biotechnol. 36, 1005.

Capecchi, M.R. (1967). Polypeptide chain termination in vitro: isolation of a release factor. Proc. Natl. Acad. Sci. U. S. A.

Capecchi, M.R., and Klein, H.A. (1970). Release factors mediating termination of complete proteins. Nature.

Chai, Y., Normam, T., Kolter, R., and Losick, R. (2009). An epigenetic switch governing daughter cell separation in Bacillus subtilis. Genes Dev. 7824, 754–765.

Chou, H.H., Delaney, N.F., Draghi, J.A., and Marx, C.J. (2014). Mapping the Fitness Landscape of Gene Expression Uncovers the Cause of Antagonism and Sign Epistasis between Adaptive Mutations. PLoS Genet. 10.

Craigen, W.J., and Caskey, C.T. (1986). Expression of peptide chain release factor 2 requires high-efficiency frameshift. Nature 322, 273–275.

Craigen, W.J., Cook, R.G., Tate, W.P., and Caskey, C.T. (1985). Bacterial peptide chain release factors: Conserved primary structure and possible frameshift regulation of release factor 2. Proc. Natl. Acad. Sci. U. S. A. 82, 3616–3620.

Dai, X., Zhu, M., Warren, M., Balakrishnan, R., Patsalo, V., Okano, H., Williamson, J.R., Fredrick, K., Wang, Y.P., and Hwa, T. (2016). Reduction of translating ribosomes enables Escherichia coli to maintain elongation rates during slow growth. Nat. Microbiol. 2, 1–9.

Dekel, E., and Alon, U. (2005). Optimality and evolutionary tuning of the expression level of a protein. Nature 436, 588–592.

Dong, H., Nilsson, L., and Kurland, C.G. (1995). Gratuitous overexpression of genes in Escherichia coli leads to growth inhibition and ribosome destruction. J. Bacteriol. 177, 1497–1504.

Duveau, F., Toubiana, W., and Wittkopp, P.J. (2017). Fitness effects of cis-regulatory variants in the saccharomyces cerevisiae TDH3 promoter. Mol. Biol. Evol. 34, 2908–2912.

Ehrenberg, M., and Kurland, C.G. (1984). Costs of accuracy determined by a maximal growth rate constraint. Q. Rev. Biophys. 17, 45–80.

Farewell, A., Kvint, K., and Nyström, T. (1998). Negative regulation by RpoS: A case of sigma factor competition. Mol. Microbiol. 29, 1039–1051.

Ferrin, M.A., and Subramaniam, A.R. (2017). Kinetic modeling predicts a stimulatory role for ribosome collisions at elongation stall sites in bacteria. Elife 6, 1–19.

Fredens, J., Wang, K., de la Torre, D., Funke, L.F.H., Robertson, W.E., Christova, Y., Chia, T., Schmied, W.H., Dunkelmann, D.L., Beránek, V., et al. (2019). Total synthesis of Escherichia coli with a recoded genome. Nature.

Gould, S.J. and R.C.L. (1978). The Spandrels of San Marco and the Panglossian Paradigm: A Critique of the Adaptationist Programme. Proc. Nat. Acad. Sci. London 205, 581–598.

Hawkins, J.S., Silvis, M.R., Koo, B., and Peters, J.M. (2019). Modulated efficacy CRISPRi reveals evolutionary conservation of essential gene expression-fitness relationships in bacteria. BioRxiv 1–15.

Hayes, C.S., and Sauer, R.T. (2003). Cleavage of the A Site mRNA Codon during Ribosome Pausing Provides a Mechanism for Translational Quality Control. Mol. Cell 12, 903–911.

Hecker, M., Pané-Farré, J., and Uwe, V. (2007). SigB-Dependent General Stress Response in *Bacillus subtilis* and Related Gram-Positive Bacteria. Annu. Rev. Microbiol. 61, 215–236.

Helmann, J.D., Wu, M., Kobel, P., Gamo, F.-J., Wilson, M., Moshedi, M., Navre, M., and Paddon, C. (2001). Global Transcriptional Response of Bacillus subtilis to Heat Shock. J. Bacteriol. 49795, 7318–7328.

Heurgué-Hamard, V., Champ, S., Engström, A., Ehrenberg, M., and Buckingham, R.H. (2002). The hemK gene in Escherichia coli encodes the N5-glutamine methyltransferase that modifies peptide release factors. EMBO J. 21, 769–778.

Hou, J., Tan, G., Fink, G.R., Andrews, B.J., and Boone, C. (2019). Complex modifier landscape underlying genetic background effects. Proc. Natl. Acad. Sci. 116, 201820915.

Hui, S., Silverman, J.M., Chen, S.S., Erickson, D.W., Basan, M., Wang, J., Hwa, T., and Williamson, J.R. (2015). Quantitative proteomic analysis reveals a simple strategy of global resource allocation in bacteria. Mol. Syst. Biol. 11, 784.

Hutchison, C.A., Chuang, R.-Y., Noskov, V.N., Assad-Garcia, N., Deerinck, T.J., Ellisman, M.H., Gill, J., Kannan, K., Karas, B.J., Ma, L., et al. (2016). Design and synthesis of a minimal bacterial genome. Science (80-.). 351, aad6253–aad6253.

Igoshin, O.A., Brody, M.S., Price, C.W., and Savageau, M.A. (2007). Distinctive Topologies of Partner-switching Signaling Networks Correlate with their Physiological Roles. J. Mol. Biol. 369, 1333–1352.

Ingolia, N.T., Ghaemmaghami, S., Newman, J.R.S., and Weissman, J.S. (2009). Genome-wide analysis in vivo of translation with nucleotide resolution using ribosome profiling. Science 324, 218–223.

Jaganathan, K., Kyriazopoulou Panagiotopoulou, S., McRae, J.F., Darbandi, S.F., Knowles, D., Li, Y.I., Kosmicki, J.A., Arbelaez, J., Cui, W., Schwartz, G.B., et al. (2019). Predicting Splicing from Primary Sequence with Deep Learning. Cell 176, 535–548.e24.

Jin, H., Björnsson, A., and Isaksson, L.A. (2002). Cis control of gene expression in E.coli by ribosome queuing at an inefficient translational stop signal. EMBO J. 21, 4357–4367.

Johnson, D.B.F., Xu, J., Shen, Z., Takimoto, J.K., Schultz, M.D., Schmitz, R.J., Xiang, Z., Ecker, J.R., Briggs, S.P., and Wang, L. (2011). RF1 knockout allows ribosomal incorporation of unnatural amino acids at multiple sites. Nat. Chem. Biol. 7, 779–786.

Johnson, D.B.F., Wang, C., Xu, J., Schultz, M.D., Schmitz, R.J., Ecker, J.R., and Wang, L. (2012). Release factor one is nonessential in escherichia coli. ACS Chem. Biol. 7, 1337–1344.

Jost, M., Santos, D.A., Saunders, R.A., Horlbeck, M.A., Hawkins, J.S., Scaria, S.M., Norman, T.M., Hussmann, J.A., Liem, C.R., Gross, C.A., et al. (2020). Titrating gene expression using libraries of systematically attenuated CRISPR guide RNAs. Nat. Biotechnol. 38, 355–364.

Karr, J.R., Sanghvi, J.C., MacKlin, D.N., Gutschow, M. V., Jacobs, J.M., Bolival, B., Assad-Garcia, N., Glass, J.I., and Covert, M.W. (2012). A whole-cell computational model predicts phenotype from genotype. Cell 150, 389–401.

Kavčič, B., Tkačik, G., and Bollenbach, T. (2019). Mechanistic origin of drug interactions between translation-inhibiting antibiotics. BioRxiv Syst. Biol.

Kearns, D.B., and Losick, R. (2005). Cell population heterogeneity during growth of Bacillus subtilis. Genes Dev. 19, 3083–3094.

Keren, L., Hausser, J., Lotan-Pompan, M., Vainberg Slutskin, I., Alisar, H., Kaminski, S., Weinberger, A., Alon, U., Milo, R., and Segal, E. (2016). Massively Parallel Interrogation of the Effects of Gene Expression Levels on Fitness. Cell 166, 1282–1294.e18.

Klumpp, S., Scott, M., Pedersen, S., and Hwa, T. (2013). Molecular crowding limits translation and cell growth. Proc. Natl. Acad. Sci. U. S. A. 110, 16754–16759.

Knöppel, A., Näsvall, J., and Andersson, D.I. (2016). Compensating the Fitness Costs of Synonymous Mutations. Mol. Biol. Evol. 33, 1461–1477.

Kurland, C.G., and Ehrenberg, M. (1987). Growth-optimizing accuracy of gene expression. Annu. Rev. Biophys. Biophys. Chem. 16, 291–317.

Lajoie, M.J., Rovner, A.J., Goodman, D.B., Aerni, H., Haimovich, A.D., Kuznetsov, G., Mercer, J. a, Wang, H.H., Carr, P. a, Mosberg, J. a, et al. (2013). Expand Biological Functions. Science 342, 357–360.

Lalanne, J.B., Taggart, J.C., Guo, M.S., Herzel, L., Schieler, A., and Li, G.W. (2018). Evolutionary Convergence of Pathway-Specific Enzyme Expression Stoichiometry. Cell 749–761.

Laursen, B., and Sørensen, H. (2005). Initiation of protein synthesis in bacteria. Microbiol. … 236, 747–771.

Li, G.W., Burkhardt, D., Gross, C., and Weissman, J.S. (2014). Quantifying absolute protein synthesis rates reveals principles underlying allocation of cellular resources. Cell 157, 624–635.

Li, X., Yokota, T., Ito, K., Nakamura, Y., and Aiba, H. (2007). Reduced action of polypeptide release factors induces mRNA cleavage and tmRNA tagging at stop codons in Escherichia coli. Mol. Microbiol. 63, 116–126.

Liu, X., Li, Y.I., and Pritchard, J.K. (2019). Trans Effects on Gene Expression Can Drive Omnigenic Inheritance. Cell 177, 1022–1034.e6.

Locke, J.C.W., Young, J.W., Fontes, M., Jiménez, M.J.H., and Elowitz, M.B. (2011). Stochastic pulse regulation in bacterial stress response. Science. 334, 366–369.

Lopatkin, A.J., and Collins, J.J. (2020). Predictive biology: modelling, understanding and harnessing microbial complexity. Nat. Rev. Microbiol.

Mangano, K., Florin, T., Shao, X., Klepacki, D., and Chelysheva, I. (2020). Genome-wide effects of the antimicrobial peptide apidaecin on translation termination.

Mitarai, N., Sneppen, K., and Pedersen, S. (2008). Ribosome Collisions and Translation Efficiency: Optimization by Codon Usage and mRNA Destabilization. J. Mol. Biol. 382, 236–245.

Mora, L., Heurgué-Hamard, V., De Zamaroczy, M., Kervestin, S., and Buckingham, R.H. (2007). Methylation of bacterial release factors RF1 and RF2 is required for normal translation termination in vivo. J. Biol. Chem. 282, 35638–35645.

Mukherjee, S., and Kearns, D.B. (2014). The Structure and Regulation of Flagella in Bacillus subtilis. Annu. Rev. Genet. 48, 319–340.

Muller, R., Meacham, Z.A., Ferguson, L., and Ingolia, N. (2020). CiBER-seq dissects genetic networks by quantitative CRISPRi profiling of expression phenotypes. BioRxiv 2020.03.29.015057.

Nakahigashi, K., Kubo, N., Narita, S.-i., Shimaoka, T., Goto, S., Oshima, T., Mori, H., Maeda, M., Wada, C., and Inokuchi, H. (2002). HemK, a class of protein methyl transferase with similarity to DNA methyl transferases, methylates polypeptide chain release factors, and hemK knockout induces defects in translational termination. Proc. Natl. Acad. Sci. 99, 1473–1478.

Ostrov, N., Beal, J., Ellis, T., Benjamin Gordon, D., Karas, B.J., Lee, H.H., Lenaghan, S.C., Schloss, J.A., Stracquadanio, G., Trefzer, A., et al. (2019). Technological challenges and milestones for writing genomes. Science. 366, 310–312.

Palmer, A.C., Chait, R., and Kishony, R. (2018). Nonoptimal Gene Expression Creates Latent Potential for Antibiotic Resistance. Mol. Biol. Evol. 35, 2669–2684.

Pané-farré, J., Quin, M.B., Lewis, R.J., and Marles-wright, J. (2017). Structure and Function of the Stressosome Signalling Hub.

Parker, D.J., Demetci, P., and Li, G.W. (2019). Rapid accumulation of motility-activating mutations in resting liquid culture of Escherichia coli. J. Bacteriol. 201, 3–6.

Parker, D.J., Lalanne, J.-B., Kimura, S., Johnson, G.E., Waldor, M.K., and Li, G.-W. (2020). Growth-Optimized Aminoacyl-tRNA Synthetase Levels Prevent Maximal tRNA Charging. Cell Syst. 1–10.

Patwardhan, R.P., Lee, C., Litvin, O., Young, D.L., Pe’Er, D., and Shendure, J. (2009). High-resolution analysis of DNA regulatory elements by synthetic saturation mutagenesis. Nat. Biotechnol. 27, 1173–1175.

Patwardhan, R.P., Hiatt, J.B., Witten, D.M., Kim, M.J., Smith, R.P., May, D., Lee, C., Andrie, J.M., Lee, S.I., Cooper, G.M., et al. (2012). Massively parallel functional dissection of mammalian enhancers in vivo. Nat. Biotechnol. 30, 265–270.

Peters, J.M., Colavin, A., Shi, H., Czarny, T.L., Larson, M.H., Wong, S., Hawkins, J.S., Lu, C.H.S., Koo, B.M., Marta, E., et al. (2016). A comprehensive, CRISPR-based functional analysis of essential genes in bacteria. Cell 165, 1493–1506.

Petersohn, A., Brigulla, M., Haas, S., Hoheisel, J.D., Völker, U., and Heckler, M. (2001). Global analysis of the general stress response of Bacillus subtilis. J. Bacteriol. 183, 5617–5631.

Petry, S., Brodersen, D.E., Murphy IV, F. V., Dunham, C.M., Selmer, M., Tarry, M.J., Kelley, A.C., and Ramakrishnan, V. (2005). Crystal structures of the ribosome in complex with release factors RF1 and RF2 bound to a cognate stop codon. Cell 123, 1255–1266.

Pollock, D.D., Thiltgen, G., and Goldstein, R.A. (2012). Amino acid coevolution induces an evolutionary Stokes shift. Proc. Natl. Acad. Sci. U. S. A. 109.

Price, C.W., Fawcett, P., Cérémonie, H., Su, N., Murphy, C.K., and Youngman, P. (2001). Genome-wide analysis of the general stress response in Bacillus subtilis. Mol. Microbiol. 41, 757–774.

Price CW (2002). General stress response. In Bacillus subtilis and Its Closest Relatives. From Genes to Cells 369–384.

Qi, L.S., Larson, M.H., Gilbert, L.A., Doudna, J.A., Weissman, J.S., Arkin, A.P., and Lim, W.A. (2013). Repurposing CRISPR as an RNA-γuided platform for sequence-specific control of gene expression. Cell 152, 1173–1183.

Rosenberg, A.B., Patwardhan, R.P., Shendure, J., and Seelig, G. (2015). Learning the Sequence Determinants of Alternative Splicing from Millions of Random Sequences. Cell 163, 698–711.

Saito, K., Green, R., and Buskirk, A.R. (2020). Translational initiation in E. coli occurs at the correct sites genome-wide in the absence of mRNA-rRNA base-pairing. Elife 9, 1–19.

Sample, P.J., Wang, B., Reid, D.W., Presnyak, V., McFadyen, I.J., Morris, D.R., and Seelig, G. (2019). Human 5′ UTR design and variant effect prediction from a massively parallel translation assay. Nat. Biotechnol. 37, 803–809.

Schaechter, M., MaalOe, O., and Kjeldgaard, N.O. (1958). Dependency on Medium and Temperature of Cell Size and Chemical Composition during Balanced Growth of Salmonella typhimurium. J. Gen. Microbiol. 19, 592–606.

Schober, A.F., Mathis, A.D., Ingle, C., Park, J.O., Chen, L., Rabinowitz, J.D., Junier, I., Rivoire, O., and Reynolds, K.A. (2019). A Two-Enzyme Adaptive Unit within Bacterial Folate Metabolism. Cell Rep. 27, 3359–3370.e7.

Scolnick, E., Tompkins, R., Caskey, T., and Nirenberg, M. (1968). Release factors differing in specificity for terminator codons. Proc. Natl. Acad. Sci. U. S. A. 61, 768–774.

Scott, M., Gunderson, C.W., Mateescu, E.M., Zhang, Z., Hwa, T., Gunderson, C.W., Mateescu, E.M., Zhang, Z., and Hwa, T. (2010). Interdependence of Cell Growth Origins and Consequences. Science. 330, 1099–1102.

Scott, M., Klumpp, S., Mateescu, E.M., and Hwa, T. (2014). Emergence of robust growth laws from optimal regulation of ribosome synthesis. 1–14.

Shah, P., McCandlish, D.M., and Plotkin, J.B. (2015). Contingency and entrenchment in protein evolution under purifying selection. Proc. Natl. Acad. Sci. U. S. A. 112, E3226–E3235.

Sharon, E., Kalma, Y., Sharp, A., Raveh-Sadka, T., Levo, M., Zeevi, D., Keren, L., Yakhini, Z., Weinberger, A., and Segal, E. (2012). Inferring gene regulatory logic from high-throughput measurements of thousands of systematically designed promoters. Nat. Biotechnol. 30, 521–530.

Shendure, J., Findlay, G.M., and Snyder, M.W. (2019). Genomic Medicine–Progress, Pitfalls, and Promise. Cell 177, 45–57.

Smith, A.M., Heisler, L.E., Mellor, J., Kaper, F., Thompson, M.J., Chee, M., Roth, F.P., Giaever, G., and Nislow, C. (2009). Quantitative phenotyping via deep barcode sequencing. Genome Res. 19, 1836–1842.

Starr, T.N., Flynn, J.M., Mishra, P., Bolon, D.N.A., and Thornton, J.W. (2018). Pervasive contingency and entrenchment in a billion years of Hsp90 evolution. Proc. Natl. Acad. Sci. U. S. A. 115, 4453–4458.

Stoebel, D.M., Dean, A.M., and Dykhuizen, D.E. (2008). The cost of expression of Escherichia coli lac operon proteins is in the process, not in the products. Genetics 178, 1653–1660.

Tubulekas, I., and Hughes, D. (1993). Growth and translation elongation rate are sensitive to the concentration of EF-Tu. Mol. Microbiol. 8, 761–770.

Urtecho, G., Insigne, K., Tripp, A.D., Brinck, M., Lubock, N.B., Kim, H., Chan, T., and Kosuri, S. (2020). Genome-wide Functional Characterization of Escherichia coli Promoters and Regulatory Elements Responsible for their Function. BioRxiv 2020.01.04.894907.

Wannier, T.M., Kunjapur, A.M., Rice, D.P., McDonald, M.J., Desai, M.M., and Church, G.M. (2018). Adaptive evolution of genomically recoded Escherichia coli. Proc. Natl. Acad. Sci. U. S. A. 115, 3090–3095.

You, C., Okano, H., Hui, S., Zhang, Z., Kim, M., Gunderson, C.W., Wang, Y.P., Lenz, P., Yan, D., and Hwa, T. (2013). Coordination of bacterial proteome with metabolism by cyclic AMP signalling. Nature 500, 301–306.

Zhu, B., and Stülke, J. (2018). SubtiWiki in 2018: From genes and proteins to functional network annotation of the model organism Bacillus subtilis. Nucleic Acids Res. 46, D743–D748.

Zhu, M., Dai, X., and Wang, Y.-P. (2016). Real time determination of bacterial in vivo ribosome translation elongation speed based on LacZα complementation system. Nucleic Acids Res. 44, gkw698.

Zhu, M., Mori, M., Hwa, T., and Dai, X. (2019). Disruption of transcription-translation coordination in Escherichia coli leads to premature transcriptional termination. Nat. Microbiol. 4, 2347–2356.

