## Supplementary material for "Spurious regulatory connections dictate the expression-fitness landscape of translation termination factors": Methods with supplementary figures

### Methods Table of Contents

#### RESOURCE AVAILABILITY

**Lead contact:**

**Materials Availability:**
Strains used in this study are available upon reasonable request.

**Data and Code Availability:**
High-throughput datasets generated in this study have been deposited to the Gene Expression Omnibus, accession GSE120861. Already published datasets (Lalanne et al., 2018) (GEO, accession GSE95211) were also used. Custom MATLAB scripts used for expression and relative fitness quantification, and data visualization are available upon reasonable request.

### EXPERIMENTAL MODEL AND SUBJECT DETAILS

**Strains**

RF inducible and control strains used for the competition experiments are listed in Supplementary Data 1. Strains used for expression profiling (RNA-seq) were GLB115 (wild-type), GLB426 (independently inducible RF2 and PrmC), GLB430 (independently inducible RF2 and PrmC, *sigB* knockout), GLB434 (control with blank ectopic constructs at *amyE* and *lacA*), GLB438 (independently inducible RF1 and PrmC), GLB442 (independently inducible RF1 and PrmC, *sigB* knockout), GLB446 (control with blank ectopic constructs at *amyE* and *levB*). Of note, our lab strain of *B. subtilis* subsp. 168 has the *swrAA*$^+$ allele (Calvio et al., 2005) (as determined by RNA-seq reads mapped to *swrAA*) leading to expression of motility gene in exponential growth, in contrast to other subsp. 168 and laboratory strains (Kearns et al., 2004).

Strains with *rsbV* UGA (wild-type), UAA and UGA variants with endogenous release factors' loci are GLB115, GLB 450, and GLB451 respectively. Strains with *rsbV* stop UGA, UAA, and UAG variants with independently inducible RF2 (xylose promoter) and RF1/PrmC/YwkF (the native RF1 operon under IPTG inducible promoter) are respectively GLB452, GLB453, and GLB454. See Supplementary Data 1

Strains for the CRISPRi experiment were a kind gift of Prof. Jason M. Peters (CAG74829: sgRNA to *prfA*, CAG74815: sgRNA to *prfB*) (Peters et al., 2016). These have sgRNA under constitutive expression ($P_{veg}$ promoter) inserted at *amyE* (with chloramphenicol resistance cassette), and catalytically inactive Cas9 under xylose promoter integrated at *lacA* (with erythromycin resistance cassette).

Strain with matched ribosome profiling and Rend-seq data for translation efficiency calibration of RF1 and RF2 expression constructs was GLB452 (see details below). Other datasets with matched ribosome profiling and Rend-seq data used solely for calibration between the transcriptome and proteome synthesis fractions for $\boldsymbol{\sigma}^B$ and translation sectors involved strains GLB372 (deletion of *ylbF*), and GLB455 (inducible green and red fluorescence proteins).

A list of plasmids and oligos used for molecular cloning can be found in Supplementary Data 2.

**Strain construction**

Standard protocol relying on natural competence of *B. subtilis* and on recombination via flanking homology sequences were used (Harwood, C R and Cutting, 1990). Molecular cloning of plasmids and recombinant DNA relied on isothermal assembly.

Two types of genetic modification were made: ectopic integration via flanking homology with resistance cassettes (Guérout-Fleury et al., 1995) (for inducible expression systems), and markerless genetic modification (clean genetic modification with no resistance marker, used for: barcode diversity generation, endogenous RF copy deletions, and *rsbV* stop codon variant generation). The pminiMAD2 strategy (single cross-over integration of modification plasmid followed by counter-selection based on a temperature sensitive origin of replication) developed by Patrick and Kearns (Patrick and Kearns, 2008) was used for markerless cloning in *B. subtilis*.

Cloning in RF inducible strains was performed with inducers appropriate for endogenous expression (25 $\mu$M IPTG, 0.035% w/v xylose) throughout growth and plating steps. In all instance of markerless cloning, full swap out of the cloning cassette was confirmed by PCR with primers outside the homology regions. All strains were confirmed by Sanger sequencing of appropriate PCR products obtained from genomic DNA of candidate clones.

RF inducible strains

As our inducible expression systems (ectopic insertion cassettes illustrated in Fig. S1b-c), we used promoters $P_{xyl}$ (with repressor XylR, derived from pDR160 (Bose and Grossman, 2011), a kind gift from Prof. Alan Grossman) responsive to xylose, and $P_{spankHy}$ (with repressor LacI, derived from pDR111 (Quisel et al., 2001), a kind gift from Prof. Rich Losick) responsive to IPTG. The $P_{xyl}$ was used without the *xylA* mini-ORF for the expressed proteins. The $P_{xyl}$ promoter has a unimodal response upon xylose induction (Peters et al., 2016). The response of LacI to IPTG is also unimodal in the absence of the lac permease LacY (Marbach and Bettenbrock, 2012) which is *de facto* not present in *B. subtilis*.

We extensively tested stability of induction of both $P_{xyl}$ and $P_{spankHy}$ promoters at different cell densities by RT-qPCR (data not shown). These confirmed that expression from these ectopic constructs were constant throughout and beyond the range ($OD_{600}$ between 0.05 to 0.4) of cell densities attained during our competition experiments and expression measurements. Projections of attained RF1, RF2, and PrmC in the 3D subspace of expression (Fig. S1e) are shown in Fig. S1f (quantified by multiplexed RNA-seq calibrated with ribosome profiling) for our inducible strains across conditions profiled, showing ability to orthogonally modulate RF levels.

To generate RF inducible strains, ectopic expression constructs with the appropriate inducible gene copies were first inserted, following by the markerless deletion (pminiMAD2 strategy) of the essential endogenous copies. The strain with independently inducible RF1 and PrmC was challenging to generatte because of high toxicity of *B. subtilis'* RF1(alone) overexpression plasmid in *E. coli* (not shown). Hence, to generate an expression construct with RF1 individually tunable, we integrated the full RF1 operon under a $P_{spankHy}$ promoter, and then deleted *prmC* and *ywkF* from both the endogenous locus and the ectopic operon. For RF2-inducible constructs, the autoregulatory frameshift in RF2 (Craigen and Caskey, 1986; Craigen et al., 1985) was removed by removing a single nucleotide (nucleotide T at position 73 in the *prfB* gene) in the ectopic construct. Once the RF related genetic modifications were completed, strains were barcoded at the *amyE* locus for competition experiments (details below).

RF1 (gene *prfA*) and RF2 (gene *prfB*) are in operons and co-transcribed with other genes. Gene *ywkF* (unknown function) is downstream and co-transcribed with *prfA* and *prmC* (gene encoding PrmC). Upon deletion of endogenous *prfA* and *prmC*, the promoter endogenous transcript was retained to preserve *ywkF* expression. Upon removal of *prmC* only, no measurable effect on the levels of the mRNAs of *prfA* and *ywkF* mRNAs were observed (e.g., RNA-seq data from strain GLB426). RF2 (gene *prfB*) is co-transcribed downstream of gene *secA*. A partial transcription terminator separates the two genes (as determined from Rend-seq (Lalanne et al., 2018)). Upon deletion of endogenous *prfB*, no effect on the mRNA level of upstream gene *secA* was detectable (e.g., RNA-seq data from strain GLB426). The gene downstream of *prfB*, *yvjA* (unknown function), which normally not co-transcribed with *prfB* (beyond low abundance transcription termination read-through mRNAs) in endogenous conditions, had increased expression in the strains with deletion of the *prfB* gene, because of an increase in the *prfB* transcription terminator readthrough (despite its sequence being preserved by our *prfB* deletion). Identifying the causes of this change in read-through the *prfB* transcription terminator following upstream gene deletion is beyond the scope of the current work.

Barcode diversity generation

We used an 8 bp region in the *amyE* (positions 328634 to 328641 on chromosome, annotation NC_000964.3, which was part of one of the homology regions used for ectopic cassette integration) as our location to introduce chromosomal barcodes. These barcodes served as a way to mark different genotypes (or redundant strains with the same genotype) in our competition experiments.

To generate different chromosomal barcodes at those positions, we created a library of "barcode swapping" plasmids. These were pminiMAD2 (Patrick and Kearns, 2008) variants with homology to *amyE* with a set of random nucleotides at the barcode positions. We generated two such plasmid libraries: one for modification in the endogenous *amyE* genotype, and one for modification in a genotype where the *amyE* region had been disrupted by an ectopic expression construct (e.g., Fig. S1b-c). Importantly, the barcode was internal to the *amyE* region, such that readout of the region was possible using identical primers for all barcoded strains (see Fig. S2b for schematics of the steps involved barcode readout). The resulting genetic changes for a successful barcode swap was the modification of only the 8 bp inside *amyE*. Following transformation of the plasmid libraries and counter-selection, clones with modified barcodes were identified by Sanger sequencing of appropriate PCR products. This strategy was employed to barcode all strains used in our competition experiments.

#### Growth conditions

All growth was performed on a tabletop orbital shaker with vigorous aeration (220 rpm) at 37°C.

Fitness landscape measurement

Growth for concurrent fitness and expression (RNA-seq) quantification (Fig. 2, 3, S3, S4) was carried out in MC complete (MCC) defined medium (Parker et al., 2020) with glycerol as the main carbon source (1% w/v). Doubling time of wild-type *B. subtilis* in this medium (as determined by manual exponential growth curve) was $21 \pm 1$ min. Inducers xylose and IPTG were added to various concentrations to modulate RFs expression levels in our inducible strains (details in Supplementary Data 3). MCC medium was conditioned to decrease the lag phase upon back dilution of cultures during competition experiments. To condition the medium, seed cultures of wild-type *B. subtilis* (GLB115) were started from freshly streaked LB plates in unconditioned MCC medium. Once pre-culture reached $\mathrm{OD}_{600} \approx 0.2$, the cultures were dilute to $\mathrm{OD}_{600} = 0.001$ in the MCC medium to be conditioned. At $\mathrm{OD}_{600} = 0.15$, medium was vacuum filtered through a coarse filter (450 nm nitrocellulose). Given prior evidence of weak pausing on asparagine/proline codons (data not shown) from ribosome profiling in conditioned medium, asparagine and proline were re-supplemented by an additional 1× their initial concentration and the resulting medium was filter-sterilized.

To minimize sources of variability, competition experiments and growth in monoculture for expression quantification were performed on the same day, with the same batch of medium, on the same shaker, from the same pre-cultures. On the morning of the day prior to experiment, the ≈20 strains involved in each experiment were streaked out (see Supplementary Data 3 for a list of strains included in each competition experiment). Later in the day, seed cultures were initiated from these fresh plates (individual growth tube for each barcoded strain) in conditioned MCC medium with inducer concentration appropriate for approximate wild-type expression of the release factors (25 $\mu$M IPTG, 0.035% w/v xylose). The next morning (day of the experiment), for competition experiments, strain pre-cultures were pooled from the overnight cultures at $1:1$ ratios. The pool was diluted to $\mathrm{OD}_{600} \approx 0.01$ in conditioned MCC medium with 25 $\mu$M IPTG and 0.035% w/v xylose until recovery to $\mathrm{OD}_{600} = 0.1$. At that point, the pool was diluted to $\mathrm{OD}_{600} = 0.004$ in 16 mL (125 mL flasks) of medium across multiple flasks corresponding to the different inducer concentrations in the experiment. Once $\mathrm{OD}_{600}$ reached 0.1 ($\approx$ 7.8 generations) in each flask, each pool was diluted into another flask (16 mL of medium) of the same medium (pre-warmed, with same inducer concentration, and shaking) to $\mathrm{OD}_{600} = 0.004$. At the time of first transfer, 8 mL of culture was harvested (in 0.8 mL of 10:1 ethanol/phenol stop solution). This first harvest constituted time 0 of the competition experiments. Subsequently, each time $\mathrm{OD}_{600}$ reached 0.1, cultures were diluted back to $\mathrm{OD}_{600} = 0.004$ and 8 mL were harvested as above. This procedure was maintained for 4 transfers (5 harvest points including time 0), or for about 30 generations. Care was taken to maintain cultures in exponential growth throughout by these frequent dilutions at low optical density ($\mathrm{OD}_{600} \leq 0.1$).

Monocultures were started in parallel to the start of the competition for expression profiling by RNA-seq. Chosen strains (one of the barcoded strains from each specified genotypes, see Supplementary Data 3) were diluted from the overnight culture (same culture as that used for pooling of competition experiments) to $\mathrm{OD}_{600} = 0.01$ in conditioned MCC medium (with 25 $\mu$M IPTG and 0.035% w/v xylose) for recovery and grown until $\mathrm{OD}_{600}$ reached 0.1. At that point, the cultures were diluted to $\mathrm{OD}_{600} = 10^{-4}$ in 25 mL of medium (125 mL flasks) in individual flasks with various inducer concentrations. 10 mL of cells were harvested (in 1 mL of 10:1 ethanol/phenol stop solution) once $\mathrm{OD}_{600} = 0.1$.

In total, three days of competition experiments were performed, with 9 inducer conditions to move along the RF1 dimension (experiment E2, see Supplementary Data 3, Fig. 2a), 9 inducers conditions to move along RF2 dimensions (experiment E1, see Supplementary Data 3, Fig. 2b), and 7+7 inducer conditions to move along the PrmC dimension (with wild-type RF2 level, and overexpression of RF2) (experiment E3, see Supplementary Data 3, Fig. 2c-d).

CRISPRi RF depletion

CRISPRi experiments (Fig. 4, S5) were carried out in LB with xylose as the inducer of the dCas9 construct (0.04% and 0.05% xylose w/v for RF1 and RF2 strains respectively). Overnight cultures (initiated from freshly streaked plates) in LB without xylose were back diluted to $\mathrm{OD}_{600} = 3 \times 10^{-4}$ in LB with xylose and were grown until $\mathrm{OD}_{600} = 0.3$.

RT-qPCR experiments

For *rsbV* stop codon switch experiments (Fig. 5), seed cultures were started from freshly streaked plates in conditioned MCC medium with inducers (25 $\mu$M IPTG, 0.035% w/v xylose). Once $\mathrm{OD}_{600}$ reached 0.1, cultures were back diluted to $\mathrm{OD}_{600} = 10^{-4}$ in conditioned MCC medium with various inducer concentrations. Cells were harvested at $\mathrm{OD}_{600} = 0.1$. Measurement of $\boldsymbol{\sigma}^{\mathrm{B}}$ activation level under lower RF2 expression (no xylose, and strain with two copies of the xylose repressor gene) were performed as above in conditioned MCC medium with 1% glycerol (no xylose, 25 $\mu$M IPTG).

The experiment to test the *rsbV* UAA variant function by ethanol stress was performed in LB. Cultures were started from freshly streaked plates, and back diluted to $\mathrm{OD}_{600} = 3 \times 10^{-4}$. Cells (wild-type GLB115, and *rsbV* UAA variant GLB450) were harvested once at $\mathrm{OD}_{600} = 0.25$ (no stress). Ethanol was then added to 4% v/v, and cells harvested 5 min later (with ethanol stress).

Matched Rend-seq/ribosome profiling datasets

Matched Rend-seq and ribosome profiling datasets used for translation efficiency calibration and conversion between transcriptome and proteome fraction were collected by diluting overnight cultures to $\mathrm{OD}_{600} = 3 \times 10^{-4}$ in the medium respective growth medium (see Supplementary Data 4 for details of strains and growth media), and harvested at $\mathrm{OD}_{600}$ between 0.15 and 0.3.

#### METHOD DETAILS

##### Gene expression measurements

Expression datasets generated and used in this work can be found in Supplementary Data 4, together with the corresponding gene-by-gene expression quantification. Oligonucleotides used for library preparation and qPCR primers are listed in Supplementary Data 2.

RNA-seq

As our primary method to quantify gene expression genome-wide, we used a previously reported RNA-seq strategy (Parker et al., 2019). Briefly, 10 mL of culture at $OD_{600} = 0.1$ were collected and mixed with 1 mL of 10:1 ethanol/phenol stop solution by rapid inversion. Cells were pelleted at 3000 rcf for 10 min at 4°C. The supernatant was decanted and cell pellets stored at -80°C until RNA extraction. RNA was extracted using the RNeasy plus kit with gDNA eliminator column (Qiagen) following manufacturer's instructions. The concentration of RNA in samples was assayed by Qubit (Thermo Fisher) from $10\times$ dilution (RNA BR). Samples were diluted to concentration of 660 ng/$\mu$L. rRNA was removed using the MICROBExpress kit (Thermo Fisher), following manufacturer's instruction but loading 2.5 $\mu$g of RNA to the reaction, using 1/8× reaction volumes, and 1 $\mu$L of capture oligo mix. Following isopropanol precipitation, samples were resuspended in 11 $\mu$L 10 mM Tris 7, and the concentration determined by Qubit (Thermo Fisher, RNA BR). For each sample, 250 ng of rRNA removed RNA was diluted in 40 $\mu$L of 10 mM Tris 7. The RNA was incubated at 95°C for 2 min and brought to ice. 4.4 $\mu$L of 10× RNA fragmentation reagents (Thermo Fisher) were added, and the mix placed at 95°C for 1 min 45 s, following which 5 $\mu$L of 10× stop solution was added. The fragmented RNA was purified with Oligo Clean and Concentrator columns (Zymo) and eluted in 17 $\mu$L 10 mM Tris 7. 3 $\mu$L of T4 PNK master mix (2 $\mu$L 10× PNK buffer, 0.5 $\mu$L SUPERase·In (Thermo Fisher), 0.5 $\mu$L T4 PNK enzyme (NEB)) was added per sample, mixed, and incubated for 60 min at 37°C, followed by 10 min at 75°C. 10 $\mu$L of PolyA master mix (3 $\mu$L 500 mM KCl, 3 $\mu$L 10 mM ATP, 2 $\mu$L 5× FS buffer (from SuperScript III enzyme), 0.5 $\mu$L SUPERase·In, 1 $\mu$L water, and 0.5 $\mu$L *E. coli* poly A polymerase (NEB)) was then added per sample, mixed, and incubated at 37°C for 30 min, followed by 10 min at 75°C. 1 $\mu$L of 25 $\mu$M indexed poly-dT reverse transcription primers (each sample having its own index, allowing pooling of samples) was added per sample, mixed, and incubated at 65°C for 5 min. The samples were returned to ice, and 9 $\mu$L of reverse transcription master mix was added per sample (3 $\mu$L 0.1 M DTT, 2 $\mu$L 10 mM dNTP mix, 2 $\mu$L 5× FS buffer (from SuperScript III enzyme), 1 $\mu$L water, 0.5 $\mu$L SUPERase·In and 0.5 $\mu$L SuperScript III (Thermo Fisher)), mixed and incubated at 50°C for 60 min, followed by 75°C for 10 min. Samples with different indices from reverse transcription primers were then pooled and the RNA hydrolyzed by adding 0.1× volume of 1M NaOH, followed by 15 min incubation at 95°C. cDNA in the range 100 to 120 nt was then size selected on a 10% TBU polyacrylamide gel (Thermo Fisher), gel extracted, and isopropanol precipitated. The cDNA was then resuspended in 20 $\mu$L of 10 mM Tris 8. 10 $\mu$L of the size selected cDNA was mixed with 5 $\mu$L 100 $\mu$M ligation adapter oDP214, 3 $\mu$L water, 5 $\mu$L 10× T4 DNA ligase buffer, 5 $\mu$L 5 M betaine, 20 $\mu$L poly(ethylene glycol) 8000, and 2 $\mu$L T4 DNA ligase (NEB), mixed thoroughly, incubated at 16°C for 10 hours. The reaction was cleaned up (Oligo Clean and Concentrator,

Zymo) and eluted in 10 $\mu$L 10 mM Tris 8. The ligated cDNA was size selected (135 to 155 nt) on a 10% TBU polyacrylamide gel, gel extracted and precipitated. Low cycle number PCR was performed with Q5 DNA polymerase (NEB) (standard reaction mix) with primers oDP007 and oDP010 (98°C for 30 s denaturation, with cycles of 10 s at 98°C, 10 s at 60°C, and 7 s at 72°C). 5 to 7 cycles of amplification were usually sufficient. The final library was size selected on 8% TBE polyacrylamide gel, gel extracted and isopropanol precipitated. Details of the final amplicon for the RNA-seq libraries can be found in Supplementary Data 4.

Sequencing data was processed as follows. Poly A tails were stripped (retaining read portion upstream of 18 A residues, for reads without 18 A residues, reads ending with at least 16 A residues were also retained), and resulting reads aligned to the *B. subtilis* genome using bowtie (option v1 k1) (Langmead et al., 2009). The 3' ends of mapped reads were summed at each genomic position. To quantify gene expression, the average read density (excluding 20 bp gaps from the annotated start and end of genes, i.e., from start+20 to end-20) was computed, and is reported in rpkm (reads per kilobase per million mapped reads) by normalizing by the total read counts not mapping to rRNA or tRNAs. Quantification for each gene in each dataset can be found in Supplementary Data 4.

mRNA levels from RF perturbed cells were compared to average mRNA levels from unperturbed datasets (average across biological replicates). Following the lack of observed changes in strains with blank expression cassettes controls (e.g., Fig. S2i-j), these datasets were also included in estimating the average unperturbed mRNA levels for each gene (see Supplementary Data 4 for list).

The current RNA-seq cDNA library preparation method compares favorably with a different library preparation approach (Rend-seq (Lalanne et al., 2018)) with different 3' and 5' adapter molecular cloning approaches: 10th and 90th percentile of fold-changes between the two methods for genes with >100 reads mapped ranged between 0.78 to 1.23 from libraries prepared starting with the same RNA material (data not shown). The above RNA-seq method is also highly reproducible, with biological replicates harvested and with libraries prepared on different days from different biological samples (same strain and growth conditions) having 10th and 90th percentile fold-changes of 0.87 and 1.13 for genes with $> 100$ mapped reads (data not shown).

<u>RT-qPCR experiments</u>

10 mL of culture were collected and mixed with 1 mL of 10:1 ethanol/phenol stop solution by rapid inversion. Cells were pelleted at 3000 rcf for 10 min at 4°C. The supernatant was decanted and cell pellets stored at -80°C until RNA extraction. RNA was extracted using the RNeasy plus kit with gDNA eliminator column (Qiagen) following manufacturer's instructions, and eluted in 10 mM Tris 7. The concentration of RNA in samples was assayed by spectrophometry (ND-1000, ThermoFisher) from a $10\times$ dilution. Samples were diluted to concentration of 1 $\mu$g/$\mu$L. 1 $\mu$L of RNA was added to 1 $\mu$L of 100 $\mu$M random hexamers, heated for 5 min at 65°C, and placed on ice. 8 $\mu$L of RT master mix (1 $\mu$L 10x reaction buffer, 0.5 $\mu$L 10 mM dNTPs mix, 0.5 $\mu$L M-MuLV reverse transcriptase (New England Biolabs), and 6 $\mu$L DEPC treated water) was added and mixed by pipetting. The mix was incubated 5 min at 25°C, 60 min at 42°C, and 20 min at 65°C. For each RNA sample, reactions with and without reverse transcriptase were run in

parallel. The RNA was subsequently hydrolyzed by adding 2 $\mu$L of 1 M NaOH and heating to 95°C for 5 min. The solution was neutralized with 2 $\mu$L of 1 M hydrochloric acid, and the reaction volume was brought to 100 $\mu$L with DEPC-treated water. For qPCR, 5 $\mu$L of Kappa SYBR green master mix, 2 $\mu$L of 1 $\mu$M forward and reverse primers, and 3 $\mu$L of diluted cDNA (above) were mixed. Reactions were monitored on a LightCycler 480 system (Roche), and Ct values obtained from the maximal second derivative method from the machine software. For each sample, multiple qPCR primer pairs were run in parallel. Samples were run on technical triplicates on 384 well plates, and large outliers among triplicates (difference in Ct value > 0.2 from technical replicates, typically less than 5% of wells) were excluded. Large differences in Ct values (>7) for reactions with and without reverse transcriptase were confirmed for each sample/primer pairs in each experiment. The mean Ct value among technical replicates was calculated and used for expression quantification. Primers to constitutively expressed genes *gyrA* were used for loading normalization. For a gene of interest, the mRNA levels relative to that of *gyrA* was calculated as $2^{Ct_{gyrA}-Ct_{oi}}$ (qPCR primer efficiencies were estimated to not differ substantially from 2 by serial dilution experiments, data not shown). Quantification from RT-qPCR experiments can be found in Supplementary Data 8.

Rend-seq/Ribosome profiling calibration

Ribosome profiling was performed as detailed previously (Lalanne et al., 2018; Li et al., 2014). Rend-seq (end enriched RNA-seq was performed as described (Lalanne et al., 2018). Given the small effect of ribosome pausing corrections and lack of 5' to 3' ramp in *B. subtilis* (Lalanne et al., 2018), relative protein synthesis rates were estimated directly as the mean ribosome footprint read density over genes (excluding 20 bp gaps from the annotated start and end of genes, i.e., from start+20 to end-20). mRNA levels from Rend-seq were quantified as the read density over gene bodies (excluding 20 bp gaps from start/end of genes as above). Rend-seq and ribosome profiling expression quantification can be found in Supplementary Data 4.

To calibrate the expression of RF1 and RF2 in our RF inducible strains, we used concurrent ribosome profiling and RNA-seq data (using Rend-seq library preparation strategy) in strain GLB452 (see Supplementary Data 1). GLB452 has RF2 without frameshift under $P_{xyl}$ and the full RF1 operon (RF1, PrmC, and YwkF) under $P_{spankHy}$. Endogenous copies of *prfA* (RF1 gene) and *prfB* (RF2 gene) are deleted. The translation efficiency (per mRNA rate of translation, denoted TE) for the exogenous copies of RF2 and RF1 were determined as ribosome profiling rpkm divided by Rend-seq rpkm, and used to derive a fold-change TE compared to wild-type mRNAs for these genes, leading to $3.0 \pm 0.4$ and $0.66 \pm 0.05$ for RF2 and RF1 respectively (error bar from standard error of the mean from 5 different RNA-seq/ribosome profiling datasets pairs at different induction levels for the constructs). Expression (measured by RNA-seq) for RF1 and RF2 in comparisons plots were corrected by the above fold-change in TE above.

Slight differences between strains used for ribosome profiling (above) and the RF inducible strains used for competition experiments are as follows. RF1 expression construct in strain GLB438 and GLB442 (used for competition experiment and fitness landscape determination) is different in that genes *prmC* and *ywkF* are not present (in frame deletion for *prmC*) from the ectopic transcript under $P_{spankHy}$. The 5' UTR and ribosome binding site for the RF1 gene are identical to those of GBL452 (in which translation efficiency calibration was performed, detailed above). We therefore use the GBL452 calibrated TE for the *prfA* mRNA in strains GLB438 and GLB442. Strains with inducible RF2 expression in competition (strains GLB426, GLB430) use the exact same construct as GLB452, with expression of RF2 coming from the same exogenous mRNA. The translation efficiency of the *prfB* mRNA calibration from GLB452 is thus directly applicable to GLB426 and GLB430. For exogenous inducible PrmC copies (under $P_{spankHy}$ in GLB426 and GLB430, and under $P_{xyl}$ for GLB438 and GLB442), we do not have corresponding TE estimates from ribosome profiling. The endogenous translation efficiency of PrmC is 0.51 (38th percentile). As a parsimonious estimate, we assume no change in the translation efficiency of our exogenous construct, which is consistent with the optimum in the expression fitness landscape corresponding to close to the expected endogenous position (Fig. 2g).

To convert our mRNA level quantification from RNA-seq to a calibrated proteome fraction (as shown on the fitness landscape plots, e.g., Fig. 2e-h,), we take the wild-type proteome fraction (estimated as the proteome synthesis fraction, see below), multiplied by the fold-change in mRNA level (directly measured by RNA-seq), and finally multiplied by the fold-change in translation efficiency (as discussed above) for the exogenous constructs.

#### Estimating the proteome mass fraction

The conversion from ribosome profiling data to proteome mass fraction relies on ribosome profiling providing an accurate estimate of protein synthesis, and on proteins being generally stable over the duration of a cell doubling time. In bacterial cells, the overwhelming majority of proteins have degradation rates small compared to the dilution arising from cell growth (Larrabee et al., 1980). For ribosome footprint density to provide an accurate measurement of protein synthesis, two assumptions need to be met: (1) the majority of ribosomes that initiate translation complete the full peptide, and (2) the average translation elongation rate is uniform across transcripts. Prior comparison to measured abundances in the literature, and assessment of protein production among stoichiometric obligatory complexes, have shown ribosome profiling to provide an accurate and precise measure of protein synthesis (Li et al., 2014). Here, the proteome synthesis fraction for a given gene was calculated from ribosome profiling data as the estimated synthesis rate multiplied by the gene size (proportional to the total number of ribosome footprint reads mapping to the gene), divided by the sum of this quantity over all genes. For a fully stable proteome, the protein synthesis fraction then equals to the proteome mass fraction.

#### Regulon transcriptome fraction

The list of annotated regulons with gene members in *B. subtilis* was downloaded from SubtiWiki (Zhu and Stülke, 2018). The transcriptome fraction for each gene was determined as the total number of RNA-seq reads mapping to member genes, and divided by the total number of reads not mapping to rRNA or tRNAs. Regulon transcriptome fraction was equal to the sum of transcriptome fraction for genes annotated as members of the regulon. The list of highlighted genes in specific regulons (Fig. 3a-b, 4b-c, S2i-j, S3a-d, S3j, S3o-p, S4c-e) can be found in Supplementary Data 5.

Estimating regulon proteome fraction

We compiled acquired datasets for which Rend-seq (mRNA level quantification) and ribosome profiling (measure of protein synthesis) were obtained from the same culture. These were used to respectively quantify the transcriptome fraction $\psi$ and proteome fraction $\phi$ (see "Estimating the proteome mass fraction" above for the conversion from ribosome profiling data to proteome fraction) for regulons of interest, here the $\boldsymbol{\sigma}^{B}$ regulon and the set of mRNA translation proteins.

For translation proteins, the relationship between proteome and transcriptome fraction was well captured by $\phi_R = \alpha\psi_R$ (with $\alpha \approx 1.1$) across our matched Rend-seq/ribosome profiling datasets, although the range of observed values for $\phi_R$ was limited (from 0.30 to 0.42). This conversion factor was used to estimate the proteome fraction from transcriptome fraction (excluding RF1, RF2, and PrmC to avoid confounding the contribution of these factors resulting from overexpression) determined by RNA-seq.

For the $\boldsymbol{\sigma}^{B}$ regulon, the relationship capturing the matched datasets was $\Delta\phi_{SigB} = \alpha\Delta\psi_{SigB}$ ($\alpha \approx 1.41$), where $\Delta\phi_{SigB} := \phi_{SigB} - \phi_{SigB}^{\circ}$ and $\Delta\psi_{SigB} := \psi_{SigB} - \psi_{SigB}^{\circ}$ are the excess proteome and transcriptome fractions from the respective basal values $\phi_{SigB}^{\circ}$ and $\psi_{SigB}^{\circ}$. Hence, we find empirically that $\phi_{SigB}^{\circ} \approx \psi_{SigB}^{\circ}$, but $\Delta\phi_{SigB} > \Delta\psi_{SigB}$. This indicates that upon induction of $\boldsymbol{\sigma}^{B}$ regulon genes, the average translation efficiency of the mRNAs of regulon genes increases. This coarse-grained observation is mechanistically corroborated, as we found numerous examples for which the production of new mRNA isoforms as a result of increased in activity in alternative promoters driven by $\boldsymbol{\sigma}^{B}$ leads to large increase in translation efficiency compared to the basal isoform expressed by housekeeping factor $\boldsymbol{\sigma}^{A}$ (McCormick and Lalanne et al, in preparation). To estimate the proteome fraction to the $\boldsymbol{\sigma}^{B}$ regulon from multiplexed RNA-seq transcriptome fraction, we use $\phi_{SigB} = \alpha\Delta\psi_{SigB} + \phi_{SigB}^{\circ}$, with $\alpha \approx 1.41$.

#### Growth rate measurement by pooled competition

We use high-throughput sequencing to quantify the relative proportion of different barcoded strains in our competition experiments, relying on the relative proportion of different barcode reads in amplicon pools as a proxy for the relative proportion of cells harboring each barcode. To readout the barcodes, we use two steps of nested PCR (Fig. S2b), appending an index encoded in primers at each step, allowing us to pool and multiplex measurement from different samples. The final amplicon is compatible with RNA-seq and ribosome profiling libraries, allowing pooling on the same sequencing lane. The approach is similar to Bar-seq (Smith et al., 2009) and derivatives, but instead of sequencing a highly complex pool at two time points, we consider a pool of limited number of barcodes ($\approx 20$ strains) at multiple time points in order to improve precision of the readout (Parker et al., 2020). Compared to previous high-precision approaches to measuring fitness based on luminescence (Kavčič et al., 2019; Kishony and Leibler, 2003) or flow cytometry (Duveau et al., 2017, 2018; Gallet et al., 2012)), our method could in principle be further improved by increasing the number of barcoded strains per genotype, which provide semi-biological replicates within each pooled experiment.

Chromosomal barcode readout

8 mL of culture (from the pool of barcoded strains in competition) was collected at $\mathrm{OD}_{600} = 0.1$ and mixed with 0.8 mL of 10:1 ethanol/phenol stop solution by rapid inversion. Cells were pelleted at 3000 rcf at 4°C for 10 min and the supernatant decanted. The cell pellets were stored at -80°C until DNA extraction. To extract genomic DNA (gDNA), we used the Promega Wizard gDNA extraction kit, following the manufacturer's protocol (scaling reagents volumes by 1/3×). The resulting gDNA was quantified on Qubit (Thermo Fisher, dsDNA BR). Samples were diluted to a concentration of 100 ng/$\mu$L. The first PCR was performed using the standard Phusion DNA polymerase (NEB) reaction to final volume of 12.5 $\mu$L, with 775 ng of template gDNA and an indexed primer with 16 nt UMI (10 variants of PCR1_UMI_FOR with different indices) and common reverse primer (oJBL124). These primers have annealing regions outside the barcodes in *amyE* common to all strains, see Fig. S2a-b. The parameters for the first PCR (PCR1, Fig. S2b) were as follows: initial denaturation at 98°C for 30 s, followed by 3 cycles of denaturation (98°C for 10 s), annealing (63°C for 30 s), elongation (72°C for 10 s). Following the first PCR, the reaction mixes were put on ice. Reactions with different indices from this PCR were pooled, cleaned up (Zymo Clean and Concentrator 5) and eluted in 100 $\mu$L 10 mM Tris 8. To get rid of residual primers and gDNA, we further applied each cleaned up pool from the first PCR to select-a-size columns (Zymo), adding 120 $\mu$L ethanol to the eluate of the first column. Final elution was made in 36 $\mu$L 10 mM Tris 8. The second PCR (PCR2, Fig. S2b) reaction was carried out with primers annealing to common regions of amplicons from the first PCR, and served to append both a second index and adapters required for sequencing on Illumina platform (Truseq primers). The PCR followed standard Phusion DNA polymerase (NEB) reaction in 12.5 $\mu$L. The parameters for the second PCR were as follows: initial denaturation at 98°C for 30 s followed by a variable number of cycles (6 to 7 cycles usually sufficient) of denaturation (98°C for 10 s), annealing (60°C for 30 s), elongation (72°C for 6 s). The final amplicon libraries (211 bp) were size selected by gel extraction on 8% TBE polyacrilamide gels (Thermo Fisher).

The structure of the final amplicon, with primers, can be found in Supplementary Data 6. Indices for the PCR reactions for our competition experiments can be found in Supplementary Data 6.

Estimation of relative fitness

Assuming no interaction between cells in a pool, and exponential growth throughout the experiment, the number of cells for strain $mut$ will grow according to $N_{mut}(t) = N_{mut}^0 2^{t/\tau_{mut}}$, where $\tau_{mut}$ is the doubling time of strain $mut$ in this particular environment. The ratio of number of cells for strain $mut$ to wild-type (WT) then obeys:

$$\log_2\left(\frac{N_{mut}(t)}{N_{WT}(t)}\right) = \log_2\left(\frac{N_{mut}^0}{N_{WT}^0}\right) + \frac{t}{\tau_{WT}}\left(\frac{\tau_{WT}}{\tau_{mut}} - 1\right).$$

We take the number of generations to be $T_{gen} := t/\tau_{WT}$, and define the relative fitness coefficient $s := \frac{\tau_{WT}}{\tau_{mut}} - 1 = \frac{\lambda_{mut}}{\lambda_{WT}} - 1$ (where $\lambda := \log(2)\tau^{-1}$ is the growth rate). We assume the number of reads corresponding to a strain barcode (after collapsing reads with the same UMI's from the primers of first PCR) $R_{mut}$ to be proportional to the number of cells corresponding to that strain in the pool (by a factor that is constant throughout the experiment), i.e., $R_{mut}(t) \propto N_{mut}(t)$, then:

$$\log_2\left(\frac{R_{mut}(t)}{R_{WT}(t)}\right) = \log_2\left(\frac{R^0_{mut}}{R^0_{WT}}\right) + s\ T_{gen}.$$

Hence, we take relative fitness (between a strain pair) as $1 + s$, where $s$ is the slope of the $\log_2$ ratio of the barcode counts as a function of number of generations. Given that most of the strains in our pool have close to wild-type growth rates, we estimate the number of generations as $-\log_2$ of the dilution factor between harvest points (e.g., $-\log_2(70\ \mu\text{L}/16\ \text{mL}) \approx 7.8$ generations for our standard dilution protocol.

UMI collapsed barcode counts (for each strain, condition, and time point) are listed in Supplementary Data 6, together with estimated $s$ for each pair of strains for each condition profiled (from the linear fit described above).

The error on the slope (as estimated by least-square), sets the precision of our fitness measurement. Assuming independent identically distributed $\log_2$ ratio measurements with standard deviation $\sigma_{\log_2 r}$, we can derive from the expression of the slope under least-square regression that the standard deviation the slope $s$, $\sigma_s^{LS}$, is (under even time sampling) for total number of generations $T_{gen}^{tot}$ and $n_t$ sampling points:

$$\sigma_s^{LS} = \sqrt{12\ \frac{n_t - 1}{n_t + 1}\ \frac{\sigma_{\log_2 r}}{\sqrt{n_t}\ T_{gen}^{tot}}} \approx \sqrt{12}\ \frac{\sigma_{\log_2 r}}{\sqrt{n_t}\ T_{gen}^{tot}},$$

where the approximation is for large $n_t$. Preliminary tests with mixing of strains at fixed ratios and technical replicate (gDNA extraction and amplicon preparation) had shown $\sigma_{\log_2 r} \approx 0.2$. Based on these, we chose to perform experiments for $\approx 30$ generations, with $n_t = 5$ samplings to have a precision of better than 1% ($\sigma_s^{LS} \approx 0.6\%$). This estimate turned out to be close to our empirical precision based on comparison of isogenic pairs (Fig. S2e).

Fig. S2c-d shows representative examples of the fitness estimation procedure for pairs of strains in a given condition (wild-type to wild-type in Fig. S2c, and wild-type to RF inducible in Fig. S2d). Importantly, redundantly barcoded strains with the same genotype provides semi-biological replicates in the same competition pool, improving our precision. For example, 4 of the 21 strain pairs are shown for redundantly barcoded WT to WT comparisons in Fig. S2c. We report the interquartile range (25th to 75th percentile) of the isogenic pairs $s$ in our main figures (e.g., error bars in Fig. 2e-h), which are typically smaller than the plotted symbol. The measured values for all pairs of strains for the comparison of fitness upon RF2 knockdown with and without *sigB* are shown in Fig. S3q.

For comparison of the relative growth rates for RF-inducible strains with and without gene *sigB*, the small fitness effect of removing the *sigB* gene in an otherwise wild-type background was subtracted. Specifically, we report $s_{RF\ inducible,\ \Delta sigB} - \langle s_{\Delta sigB}\rangle$ , with $\langle s_{\Delta sigB}\rangle$ the average fitness defect of the *sigB* deletion across all conditions in a given competition experiment (there were slight variations in $\langle s_{\Delta sigB}\rangle$ from one competition experiment to another).

**Quality control of growth measurement**
Numerous experiments were performed to assess precision and accuracy of the barcode readout protocol and fitness measurement, as well as biological impact, independent of RF perturbations, of our ectopic expression cassettes and resistance markers.

Barcode readout cross-talk assessment
The two-step nested PCR protocol to generate our amplicon library allows us to append two sets of indices on our samples, permitting multiplexing amplicon sequencing of various conditions. Such pooling of samples prior to PCR2 can introduce cross-talk between indices and barcodes. To characterize such cross-talk, we spiked-in genomic DNA extracted from individual barcoded strains not present in the sequenced pool at specific combinations of PCR indices (Supplementary Data 6, 7). Bleed through of these barcodes to different PCR indices provides a measure of cross-talk arising (for example) from carry over indexed primers from the first PCR to the second PCR.

Supplementary Data 7 shows the results for barcode cross-talk for competition experiments. Rows correspond to indices from the first PCR (PCR1), and columns to indices from the second PCR (PCR2). PCR1 indexed samples are pooled prior to PCR2. The tables show the number reads mapping to the barcode for the spiked-in genomic DNA (each table corresponds to a different spike-in) with indices from PCR1 and PCR2. Highlighted positions in the table correspond to positions were corresponding spike-in was added. Read counts at non-highlighted positions correspond to cross-talk.

Two types of cross-talk can be distinguished: (1) intra PCR2 pool (appearance of spiked-in barcode at different PCR1 indices inside pools were the spike-in was added) corresponding to reads not highlighted in a column with a highlighted cell, and (2) inter PCR2 pool (appearance of spiked-in barcode at PCR1 indices in pools were the spike-in was not added) corresponding to reads in a column with no highlighted cell. Interpool cross-talk was very low, with relative number of reads incorrectly mapping per spike-in barcode being $< 5 \times 10^{-6}$. Intrapool cross-talk was higher, but $> 99\%$ (and typically higher) of spike-in barcode reads mapped to the correct pair of PCR indices. Prior experiments without the additional primer clean up before the select-a-size column purification after PCR1 suggested higher intrapool cross-talk, reaching $> 10\%$ in some cases (data not shown), possibly from carry-over primers from PCR1. The additional clean up step strongly reduced barcode cross-talk.

Accuracy of barcode readout
Mixing of two strains with different at predetermined ratios followed by our barcode readout procedure recovered the expected ratios across four orders of magnitudes (Fig. S2h).

Accuracy of fitness measurement
The accuracy of the method was determined by comparing the measured $s$ from competition experiment with the growth rate measured via a "manual" growth curve (measurement of optical density versus time) from the monoculture from which cells were harvested for multiplexed RNA-seq. Supplementary Data 7 lists the comparisons. Conditions for which $s$ from competition

experiment were $|s| > 0.2$ (large growth defect) and with 3 or more $\mathrm{OD}_{600}$ data points within the range 0.005 to 0.12 (reliability of manual doubling time estimate) were retained for comparison. The inferred $s$ from the growth curve was calculated as $s := \frac{\tau_{WT}}{\tau_{mut}} - 1$, with $\tau_{WT} = 21 \pm 1$ min and $\tau_{WT}$ from the linear fit of log transformed $\mathrm{OD}_{600}$ data points. Range for the doubling time measurements are estimated from bootstrap subsampled slopes. Overall, most measurements agreed within error, with the manual measurements being much less precise, and with agreement being better for larger growth defect (as expected given the noise in the manually estimated $s$ for small growth defect).

Precision of fitness measurement

A stringent measure of precision of our relative growth measurement from competition is the distribution of measured relative fitness $s$ arising from isogenic strains (apart from having a different chromosomal barcodes) across our conditions. Given that most genotypes are redundantly barcoded at least 4 times (except *ΔsigB*, with two barcodes), this provides us with 6 pairwise comparisons per condition per genotype pairs. Under the assumption that the identity of the 8 bp barcode does not affect fitness, $s$ should be 0 for these strain pairs. We emphasize that in addition to noise in the readout, non-zero $s$ could come from accumulated deleterious/beneficial mutations in the course of the barcoding cloning procedure and pre-cultures prior to competition experiments. Hence, the distribution of $s$ for isogenic pairs (examples of such isogenic pair competitions are shown in Fig. S2c) corresponds to a lower bound on the precision of the measurement readout itself. Across n=1253 such pairwise comparisons spanning all inducer conditions and experiments, we find $\sigma_s = 0.6\%$ (median $|s| = 0.3\%$), with the full distribution of isogenic $s$ (Fig. S2e), shown as inset in Fig. 2h. We take the resolution of our fitness measurement to be $\pm 2\sigma_s = 1.2\%$ (grey shading in Fig. S2e).

Impact of ectopic expression cassettes

Our RF expression constructs involve the disruption of endogenous loci (*amyE, lacA*, and *levB*), together with the addition of exogenous genes (various resistance cassettes and repressors) which could contribute to fitness defect either through cost of expression or the specific activity of these genes (although none of the resistance cassettes used in our inducible RF strains modify endogenous cell machinery). We constructed and redundantly barcoded control strains with "blank" ectopic expression cassettes, which were identical to the cassettes used to drive inducible expression of RFs, but without any genes under the inducible promoters. These control strains were included in all our competition experiments. Two suites of such control strains were made: (1) with blank insertions at *amyE* and *lacA* (GLB434-437 series), and (2) with blank insertions at *amyE* and *levB* (GLB446-449 series). See strain details in Supplementary Data 1.

Both transcriptome characterization (representative comparisons to wild-type in Fig. S2i for GLB434, and Fig. S2j for GLB446 showing little genome-wide changes) and fitness measurements (Fig. S2f-g shows the distribution of $s$ for these strains compared to wild-type across our experimental conditions) showed little impact of these ectopic cassettes. In particular, we did not see trend with $s$ as a function of increasing inducer concentration. Overall, our control strains showed growth defects of about 1% or smaller, close to the resolution of our measurement, suggesting little fitness defects arising from the disruptions to the genome and expression of exogenous proteins (resistance and repressor proteins). Given these small fitness defects for control strains, we compared our RF inducible strains directly to wild-type because of the higher number of barcoded wild-type strains, improving our precision.

Assessing the impact of the lag phase

Our competition experiments relied on periodic dilutions of cultures to maintain approximate steady-state exponential growth, as opposed to using an automated continuous culture device such as a turbidostat (Schober et al., 2019; Toprak et al., 2013; Wong et al., 2018). While conditioning our growth medium reduced lags between dilutions, and dilutions were made at the low $\mathrm{OD}_{600}$ of 0.1, a short lag was still observed at each dilution. To confirm that the observed growth defects originated from the exponential growth rate differences as opposed to slight differences in lag time between strains, we performed two series of competition experiments with the same number of total generations, but different number of dilutions (few large dilutions vs. many small dilutions). The two different dilution scenarios consisted in the same total number of generations of growth, but additional lag periods for the scenario with many small dilutions. Specifically, the first scenario consisted in 4 dilutions of 1000× (40 generations total) with harvesting at each dilution and at t=0 (scenario large dilutions $L$), and the second scenario consisted in 8 dilutions of 31.6× (40 generations total) with harvesting at t=0 and every other two dilutions (scenario small dilutions $S$). These experiments were performed with a single barcoded strain compared to wild-type (one barcode), and was with a different RF inducible system (GLB452, same as $rsbV$ stop codon switching and ribosome profiling calibration), where RF2 was under $\mathrm{P}_{xyl}$ and the full RF1 operon (RF1, PrmC, and YwkF) was under $\mathrm{P}_{spankHy}$. 9 different conditions with these two dilution protocols were considered, spanning different regions of the (RF1, RF2, PrmC) expression space. Results for measured $s$ for the two different dilution scenarios, compiled in Supplementary Data 7, showed near complete agreement between the two dilution scenarios, supporting that possible slight differences in recovery from the lag phase upon RF perturbation did not contribute to the measured growth defects, even for large perturbations.

#### Details on the $\boldsymbol{\sigma}^{\mathrm{B}}$ regulon

Selection of reporter genes in $\boldsymbol{\sigma}^{\mathrm{B}}$ regulon

To select reporter genes to assess $\boldsymbol{\sigma}^{\mathrm{B}}$ regulon activity by reverse transcription quantitative PCR (RT-qPCR), we identified a list of genes annotated as members of the $\boldsymbol{\sigma}^{\mathrm{B}}$ regulon which were consistently most highly induced transcriptionally in our RNA-seq datasets. *gsiB* was the most strongly induced $\boldsymbol{\sigma}^{\mathrm{B}}$ gene, but its sequence contained multiple duplicated regions, making its quantification by RT-qPCR unreliable. $ywzA$ and $ygxB$ were two highly and consistently induced genes in the regulon (highlighted dark red in Fig. 3a). These genes were selected as targets to monitor induction of the regulon for the $rsbV$ stop codon switching experiment.

Bioinformatic analysis of $\boldsymbol{\sigma}^{\mathrm{B}}$ operons

To search for other $sigB$ operons in diverse species, we scanned the representative and reference genomes from the RefSeq database (Tatusova et al., 2016) with protein blast (evalue cutoff

$10^{-7}$) to the three amino acid sequences of RsbV, RsbW, and $\boldsymbol{\sigma}^{B}$ from *B. subtilis*. We retained species with hits to all three proteins, extracted positions of homologous proteins on chromosomes, and identified connected clusters of hits based on a permissive spatial cutoff of 100 bp (distance between start to end of genes). Retaining connected spatial clusters with the three proteins, and also filtering based on conserved gene order and orientation, led to 95 final candidate $\boldsymbol{\sigma}^{B}$ operons (the operon with $\boldsymbol{\sigma}^{B}$ homolog with highest identity to *B. subtilis*' $\boldsymbol{\sigma}^{B}$ was retained in species with multiple candidate operons). Candidates are listed in Supplementary Data 9. Final candidates are summarized in Fig. S6. For these *sigB* operon candidates: 18/95=19% had the **AUGA** overlap between *rsbV* and *rsbW*. In comparison, only 7% of co-directional gene pairs (226/3042) have that arrangement in *B. subtilis* (Fig. S5j). Further, 42/95 = 44% of *sigB* operon candidates had any form of coding sequence overlap between *rsbV* and *rsbW*, compared to 17% (521/3042) across co-directional gene pairs in *B. subtilis*.

#### CRISPRi depletion of RF1/PrmC and RF2

As an orthogonal system to perturb expression of RF, we performed experiments in CRISPRi (dCas9 targeted to a specific locus by sgRNA, blocking transcription and decreasing gene expression by up to 100×). Strains targeting *prfB* (strain CAG74815) and *prfA* (strain CAG74829) (Peters et al., 2016), a kind gift of Prof. Jason M. Peters, were used. Given the operon structure of RF1 and PrmC (co-transcribed), the knockdown of RF1 also lead to knockdown in PrmC.

Characterization of the induction property of the xylose promoter (driving dCas9 expression in the CRISPRi strains) in LB using RT-qPCR revealed around 20× induction at 0.04% w/v xylose as the $\mathrm{OD}_{600}$ of the culture increased from 0.1 to 0.3 (data not shown). The physiological causes of such induction at intermediate $\mathrm{OD}_{600}$ are beyond the scope of the current work, but could be due to consumption of residual glucose present in LB, alleviating competitive inhibition of the xylose repressor (Dahl et al., 1995). This transient induction was used to generate an acute, but non steady-state, knockdown of RF1/PrmC and RF2. Following dilution to $\mathrm{OD}_{600} = 3 \times 10^{-4}$ in LB with xylose (0.04% for RF1, 0.05% for RF2), and cells were harvested for Rend-seq and ribosome profiling (flash filtration) at $\mathrm{OD}_{600} = 0.3$.

Quantification of mRNA levels and protein synthesis in these conditions revealed over 60× knockdown for both RF1 and RF2 (RF2 mRNA level quantified by average read density after the position targeted by the sgRNA: positions 3627150 to 3628026 in NC_000964.3), Fig. 4b-c. Together with these RF specific perturbations, our protein synthesis measurements showed large scale induction of $\boldsymbol{\sigma}^{B}$ upon RF2 knockdown (Fig. 4c), and mild *eps* gene induction in RF1/PrmC knockdown (median fold-change increase of 2.9 compared to wild-type, not highlighted in Fig. 4b), consistent with remodeling in the gene expression program observed in our steady-state measurements with our RF inducible strains.

While RF abundances are challenging to quantify from synthesis rates in non steady-state conditions, queuing was still observed at RF-specific stop codons by ribosome profiling (Fig. 4a, S5a-c), suggesting high depletion in RF. Metagene stop codon traces (Fig. 4a) were obtained as follows: for each stop codon (UAA, UAG, UGA), all genes expressed to at least 0.5 ribosome footprint read/nt were retained and their mapped footprint reads (center-mapped (Li et al., 2014)) traces (normalized by the mean density in the first half of the gene) were aligned by the stop codon. The median at each position over all such aligned traces constituted metagene plot (Fig. 4a). Each stop codons show elevated ribosome density in wild-type. However, for RF1/PrmC and RF2 knockdown, elevated densities following periodic patterns of $\approx 25$ nt (ribosome footprint size) appeared upstream of stop codons specific to the knocked down RF, Fig. 4a (bottom two rows), likely corresponding to ribosome queues forming as a result of slow translation termination. All ribosome footprint traces for genes meeting the read density threshold were also visualized as a heat map, Fig. S5a-c. More prominent queues were observed for genes with higher translation efficiency TE (as determined from wild-type data from (Lalanne et al., 2018), TE being proportional the rate of ribosome initiating on individual mRNAs), consistent with a simple stochastic theory of queue formation (schematic Fig. S5h) (Bergmann and Lodish, 1979; Mitarai et al., 2008), Supplementary Discussion.

The ribosome profiling data under acute RF knockdown was used to assess expression stoichiometry of co-directional gene pairs (Fig. 4d). To do so, the ribosome footprint density ratio for selected gene pairs was calculated in wild-type and under RF knockdown. Gene pairs, restricted to pairs with >10 reads mapping to both genes in the two conditions, were then stratified based on stop codon identity and distance between gene pairs. The distribution of fold-change of the expression stoichiometry (down/up ribosome footprint density) under RF perturbation vs. WT were then recorded under different stratifications. Fig. 4d shows the fold-change distribution for co-directional gene separated by less than 30 bp, and stratified by the stop codon of the upstream gene. It shows a modest but clear decrease in the expression of genes downstream of stop codons cognate to the RF perturbation. Additional control analyses were performed to assess the validity of this effect. First, gene pairs within 30 bp but stratified by the stop codon of the downstream gene did not show this effect, Fig. S4d. In addition, gene pairs separated by more than 30 bp stratified by the upstream stop codon did not show this effect, Fig. S4e. Performing the same analysis (stratified by upstream stop codon, within 30 bp) but from Rend-seq data (same experiment) also did not show this effect (Fig. S4f), suggesting that the decrease in expression came from changes in translation initiation, and not mRNA levels. Analyses considering fold-changes in gene expression (compared to wild-type, normalized by the median fold-changes of all genes with sufficient coverage, >10 reads mapped in both conditions, irrespective of stop codons) stratified by stop codon (Fig. 4e for ribosome profiling, Fig. S5g for mRNA levels) did not show substantial stop codon specific effects.

#### QUANTIFICATION AND STATISTICAL ANALYSIS

Gene expression was quantified from sequencing data as described in Methods Details sections.

To assess significance of the stop codon specific trends observed in our expression datasets for acute RF depletion by CRISPRi (Fig. 4d-e S4d-g), we generated random reshufflings of stop codon categories (i.e., the identities of the gene's stop codon were randomly permuted). For each stop codon reshuffling, we calculated the median of fold-change for the (reshuffled) RF-specific stop cognate to the RF perturbation and the median of the fold-change for the (reshuffled) RF-agnostic stop UAA. The ratio of these median fold-changes constituted the effect size. The p-values was taken as the fraction of stop codon reshuffling for which the effect size was more pronounced (smaller ratio of fold-changes) than the non-reshuffled list.

To assess the significance of the fitness rescue upon *sigB* deletion for the maximum steady-state knockdown of RF2 (Fig. S3q), we randomly sampled (with replacement) 12 (out of total 96) pairwise fitness differences at the non maximal RF2 knockdown. The median of these sampling was compared to the median of the sampled (12 samplings with replacement of the 12 pairwise fitness values) fitness differences under maximal RF2 knockdown. The p-value was taken as the fraction of random samplings for which the median of the fitness difference for non maximal RF2 knockdown exceeded the median of the fitness difference of maximal RF2 knockdown.

### SUPPLEMENTARY DISCUSSION

**Stochastic queuing model of ribosomes**

The number of ribosomes on an mRNA can be approximated by solving a totally asymmetric exclusion process (Shaw et al., 2003), but a further simplified model disregarding spatial information recapitulates the statistics of queue formation (as verified by full stochastic simulations, data not shown). The state space of the simplified queue model is the number of ribosomes $N$ in the queue upstream of the stop codon (where $N = 1$ when one ribosome is on the stop codon). Ribosomes arrive at a rate $\alpha$ (initiation rate on the transcript), and leave at the termination rate $\beta$. The ribosome arrival rate at the queue is rigorously equal to $\alpha$ in steady-state (we neglect mRNA degradation in this discussion), unless the queue becomes large enough to affect the initiation process (fully jammed transcript). This jammed regime is far from considered conditions, even with severe RF depletion (as determined from the typical queue length from our metagene analysis, Fig. 4a)

The stochastic process away from the jammed state is then described by: $N \rightarrow N + 1$ at rate $\alpha$ (ribosome arriving at the stop codon), and $N \rightarrow N - 1$ at rate $\beta$ (ribosome leaving the stop codon) for $N > 0$. It assumes that the processes of initiation and termination are independent of the number of ribosomes in the queue. The probability for the queue to have $N$ ribosomes, $P(N)$, can be obtained by solving the steady-state solution of the resulting master equation, leading to a geometric series: $P(N) = (\alpha/\beta)^N(1 - \alpha/\beta)$. Hence, the prevalence of higher order queues scales as the ratio of the initiation to termination rate on the transcript to the power of the number of ribosomes queued. The average queue size, i.e., $\langle N \rangle$, is $\alpha/(\beta - \alpha)$. The divergence of queue length as initiation rate $\alpha$ approaches termination rate $\beta$ corresponds to the jamming transition, with long queues. This relationship shows that upon decrease of termination rate (e.g., from RF depletion), mRNAs with larger initiation rate (translation efficiencies), are expect to have longer queues (schematically illustrated in Fig. S5h), which is what we observe on individual genes (Fig. S5b-c).

**Predictions of bacterial growth laws for gratuitous protein expression**

The *E. coli* bacterial growth laws (Box 1) formulate predictions of the impact of gratuitous expression on the translation sector compression and growth. Coarse-grained regulatory parameters ($\phi_{\circ}, \phi_R^{max}, \kappa_n, \kappa_t$) required for the predictions can be obtained from fits of physiological trajectories in the space of growth rate vs. translation sector space (Fig. 2j-l). The resulting predictions (Box 1, Eq. 1 and 2) with parameters obtained here are shown as dashed lines in Fig. 2j-l.

Central to the *E. coli* growth law predictions is the incompressibility of a large portion (Q sector in (Scott et al., 2010)) of the proteome, which leads to steeper growth defect upon gratuitous expression. In contrast, in our comparison with the estimated proteome fraction occupied by the $\boldsymbol{\sigma}^{\mathrm{B}}$ regulon, we find that growth defect is well explained by a purely passive compression factor of $1 - \phi_{\mathrm{U}}$ (no incompressible sector, full lines in Fig. 3e, 3h, S3g, S3i), c.f. Box. 1 Eq. 1 and 2. Incompressibility was hypothesized to arise from autoregulatory mechanisms, which could be species specific and not be present in *B. subtilis*.

**Tuning PrmC under RF2 overexpression**

In order to verify that the growth defect resulting from RF2 overexpression (s ≈ −6%, Fig. 2f) was not caused by imbalance between RF and PrmC, we tuned PrmC over the range achievable with our inducible construct under maximal RF2 overexpression (Fig. 2d, 2h). Increased PrmC expression above wild-type level did not measurably rescue growth, suggesting that the growth defect for RF2 overexpression was not due to lack of post-translational modification of RF2 resulting from insufficient PrmC levels.

Consistently with the observations for wild-type RF2 levels, over-expressing PrmC led to a large induction in the $\boldsymbol{\sigma}^B$ regulon (Fig. S3c), with associated translation sector compression (Fig. S3g), and growth defect (Fig. S3h). Transcriptomic re-arrangements following PrmC overexpression were highly reproducible and largely independent of RF2 levels (Fig. S3j).

In contrast to the lack of change upon PrmC overexpression, the growth defect observed for PrmC knockdown was aggravated by RF2 overexpression (Fig. S3m-n), possibly due to increased stoichiometric imbalance between PrmC and the total RF1 and RF2 concentration. Although mild $\boldsymbol{\sigma}^B$ induction was observed in these conditions (Fig. S3a, S3e), this induction was not causative of the observed growth defect, as deleting *sigB* did not rescue growth (blue shading, Fig. S3e-f, S3h) in this region of the RF expression subspace. In particular, the translation sector was compressed, seemingly because of large-scale transcriptomic changes independent of *sigB* (Fig. S3a-b and insets), in contrast to the situation for overexpression of PrmC. The identification of the events leading to these transcriptional changes will warrant further studies.

**RF1 knockdown and global changes in expression**

Upon steady-state RF1 knockdown, and associated with the sharp decrease in cell growth (Fig. 2e, S4b), we observe large changes in specific regulons (some examples shown in Fig. S4c), analogously to $\boldsymbol{\sigma}^B$ induction upon PrmC overexpression. For example, members of the motility regulon (driven by alternative sigma factor $\boldsymbol{\sigma}^D$, gene *sigD*) and *lyt* operon, have a 5× or larger median decrease in mRNA levels (Fig. S4d). mRNA levels of biofilm production matrix genes (*eps* operon) are up about 10× (Fig. S4d). These effects are independent of *sigB* (Fig. S4e).

Attempting to identify causative molecular events responsible for these changes is complicated by the large number of regulators controlling motility in *B. subtilis* (Mukherjee and Kearns, 2014). Our CRISPRi depletion of RF1/PrmC, which was not in steady-state of growth, indicates that induction of *eps* genes are likely initial steps in the cascade, as their expression is up before large changes to $\boldsymbol{\sigma}^D$ and *lyt* operons (median fold-change increase of 2.9 compared to wild-type for the data in Fig. 4b, not highlighted). Multiple regulators are known to modulate expression of *eps* genes (e.g., RemA (Winkelman et al., 2013), SinR (Kearns et al., 2005), and DegU (Murray et al., 2009)), and many have connections to RF1. *remA* is an operon with an upstream gene with the RF1 UAG stop codon, although the two genes are separated by a large distance (77 bp) and show no changes in expression stoichiometry upon RF1/PrmC depletion in our dataset (not shown). Both *sinR* and *degU* end with the RF1 UAG stop. These provide plausible candidate stop codons to modify for attempting to rewire of the RF1 landscape, analogously to our modifications of the $\boldsymbol{\sigma}^B$ operon (Fig. 5), although identification of the regulatory network's focal point in this instance lies beyond the scope of the current work.

**Methods References**

Bergmann, J.E., and Lodish, H.F. (1979). A Kinetic Model of Protein Synthesis. *254*, 11927–11937.

Bose, B., and Grossman, A.D. (2011). Regulation of horizontal gene transfer in Bacillus subtilis by activation of a conserved site-specific protease. J. Bacteriol. *193*, 22–29.

Calvio, C., Celandroni, F., Ghelardi, E., Amati, G., Salvetti, S., Ceciliani, F., Galizzi, A., and Senesi, S. (2005). Swarming differentiation and swimming motility in Bacillus subtilis are controlled by swrA, a newly identified dicistronic operon. J. Bacteriol. *187*, 5356–5366.

Craigen, W.J., and Caskey, C.T. (1986). Expression of peptide chain release factor 2 requires high-efficiency frameshift. Nature *322*, 273–275.

Craigen, W.J., Cook, R.G., Tate, W.P., and Caskey, C.T. (1985). Bacterial peptide chain release factors: Conserved primary structure and possible frameshift regulation of release factor 2. Proc. Natl. Acad. Sci. U. S. A. *82*, 3616–3620.

Dahl, M.K., Schmiedel, D., and Hillen, W. (1995). Glucose and glucose-6-phosphate interaction with Xyl repressor proteins from Bacillus spp. may contribute to regulation of xylose utilization. J. Bacteriol. *177*, 5467–5472.

Duveau, F., Toubiana, W., and Wittkopp, P.J. (2017). Fitness effects of cis-regulatory variants in the saccharomyces cerevisiae TDH3 promoter. Mol. Biol. Evol. *34*, 2908–2912.

Duveau, F., Hodgins-Davis, A., Metzger, B.P.H., Yang, B., Tryban, S., Walker, E.A., Lybrook, T., and Wittkopp, P.J. (2018). Fitness effects of altering gene expression noise in saccharomyces cerevisiae. Elife *7*, 1–33.

Gallet, R., Cooper, T.F., Elena, S.F., and Lenormand, T. (2012). Measuring selection coefficients below 10-3: Method, Questions, and Prospects. Genetics *190*, 175–186.

Guérout-Fleury, A.M., Shazand, K., Frandsen, N., and Stragier, P. (1995). Antibiotic-resistance cassettes for Bacillus subtilis. Gene *167*, 335–336.

Harwood, C R and Cutting, S.M. (1990). Molecular Biological methods for Bacillus (John Wiley).

Kavčič, B., Tkačik, G., and Bollenbach, T. (2019). Mechanistic origin of drug interactions between translation-inhibiting antibiotics. BioRxiv Syst. Biol.

Kearns, D.B., Chu, F., Rudner, R., and Losick, R. (2004). Genes governing swarming in Bacillus subtilis and evidence for a phase variation mechanism controlling surface motility. Mol. Microbiol. *52*, 357–369.

Kearns, D.B., Chu, F., Branda, S.S., Kolter, R., and Losick, R. (2005). A master regulator for biofilm formation by Bacillus subtilis. Mol. Microbiol. *55*, 739–749.

Kishony, R., and Leibler, S. (2003). Environmental stresses can alleviate the average deleterious effect of mutations. J. Biol. *2*, 14.

Lalanne, J.B., Taggart, J.C., Guo, M.S., Herzel, L., Schieler, A., and Li, G.W. (2018). Evolutionary Convergence of Pathway-Specific Enzyme Expression Stoichiometry. Cell 749–761.

Langmead, B., Trapnell, C., Pop, M., and Salzberg, S. (2009). Ultrafast and memory-efficient alignment of short DNA sequences to the human genome. Genome Biol. *10*, R25.

Larrabee, K.L., Phillips, J.O., Williams, G.J., and Larrabee, A.R. (1980). The relative rates of protein synthesis and degradation in a growing culture of Escherichia coli. J. Biol. Chem. *255*, 4125–4130.

Li, G.W., Burkhardt, D., Gross, C., and Weissman, J.S. (2014). Quantifying absolute protein synthesis rates reveals principles underlying allocation of cellular resources. Cell *157*, 624–635.

Marbach, A., and Bettenbrock, K. (2012). Lac operon induction in Escherichia coli: Systematic comparison of IPTG and TMG induction and influence of the transacetylase LacA. J. Biotechnol. *157*, 82–88.

Mitarai, N., Sneppen, K., and Pedersen, S. (2008). Ribosome Collisions and Translation Efficiency: Optimization by Codon Usage and mRNA Destabilization. J. Mol. Biol. *382*, 236–245.

Mukherjee, S., and Kearns, D.B. (2014). The Structure and Regulation of Flagella in Bacillus subtilis . Annu. Rev. Genet. *48*, 319–340.

Murray, E.J., Kiley, T.B., and Stanley-Wall, N.R. (2009). A pivotal role for the response regulator DegU in controlling multicellular behaviour. Microbiology *155*, 1–8.

Parker, D.J., Demetci, P., and Li, G.W. (2019). Rapid accumulation of motility-activating mutations in resting liquid culture of Escherichia coli. J. Bacteriol. *201*, 3–6.

Parker, D.J., Lalanne, J.-B., Kimura, S., Johnson, G.E., Waldor, M.K., and Li, G.-W. (2020). Growth-Optimized Aminoacyl-tRNA Synthetase Levels Prevent Maximal tRNA Charging. Cell Syst. 1–10.

Patrick, J.E., and Kearns, D.B. (2008). MinJ (YvjD) is a topological determinant of cell division in Bacillus subtilis. Mol. Microbiol. *70*, 1166–1179.

Peters, J.M., Colavin, A., Shi, H., Czarny, T.L., Larson, M.H., Wong, S., Hawkins, J.S., Lu, C.H.S., Koo, B.M., Marta, E., et al. (2016). A comprehensive, CRISPR-based functional analysis of essential genes in bacteria. Cell *165*, 1493–1506.

Quisel, J.D., Burkholder, W.F., and Grossman, A.D. (2001). In vivo effects of sporulation kinases on mutant Spo0A proteins in bacillus subtilis. J. Bacteriol. *183*, 6573–6578.

Schober, A.F., Mathis, A.D., Ingle, C., Park, J.O., Chen, L., Rabinowitz, J.D., Junier, I., Rivoire, O., and Reynolds, K.A. (2019). A Two-Enzyme Adaptive Unit within Bacterial Folate Metabolism. Cell Rep. *27*, 3359-3370.e7.

Scott, M., Gunderson, C.W., Mateescu, E.M., Zhang, Z., and Hwa, T. (2010). Interdependence of cell growth and gene expression: origins and consequences. Science *330*, 1099–1102.

Shaw, L.B., Zia, R.K.P., and Lee, K.H. (2003). Totally asymmetric exclusion process with extended objects: a model for protein synthesis. Phys. Rev. E. Stat. Nonlin. Soft Matter Phys. *68*, 021910.

Smith, A.M., Heisler, L.E., Mellor, J., Kaper, F., Thompson, M.J., Chee, M., Roth, F.P., Giaever, G., and Nislow, C. (2009). Quantitative phenotyping via deep barcode sequencing. Genome Res. *19*, 1836–1842.

Tatusova, T., Dicuccio, M., Badretdin, A., Chetvernin, V., Nawrocki, E.P., Zaslavsky, L., Lomsadze, A., Pruitt, K.D., Borodovsky, M., and Ostell, J. (2016). NCBI prokaryotic genome annotation pipeline. Nucleic Acids Res.

Toprak, E., Veres, A., Yildiz, S., Pedraza, J.M., Chait, R., Paulsson, J., and Kishony, R. (2013). Building a morbidostat: an automated continuous-culture device for studying bacterial drug resistance under dynamically sustained drug inhibition. Nat. Protoc. *8*, 555–567.

Winkelman, J.T., Bree, A.C., Bate, A.R., Eichenberger, P., Gourse, R.L., and Kearns, D.B. (2013). RemA is a DNA-binding protein that activates biofilm matrix gene expression in Bacillus subtilis. Mol. Microbiol. *88*, 984–997.

Wong, B.G., Mancuso, C.P., Kiriakov, S., Bashor, C.J., and Khalil, A.S. (2018). Precise, automated control of conditions for high-throughput growth of yeast and bacteria with eVOLVER. Nat. Biotechnol. *36*, 614–623.

Zhu, B., and Stülke, J. (2018). SubtiWiki in 2018: From genes and proteins to functional network annotation of the model organism Bacillus subtilis. Nucleic Acids Res. *46*, D743–D748.

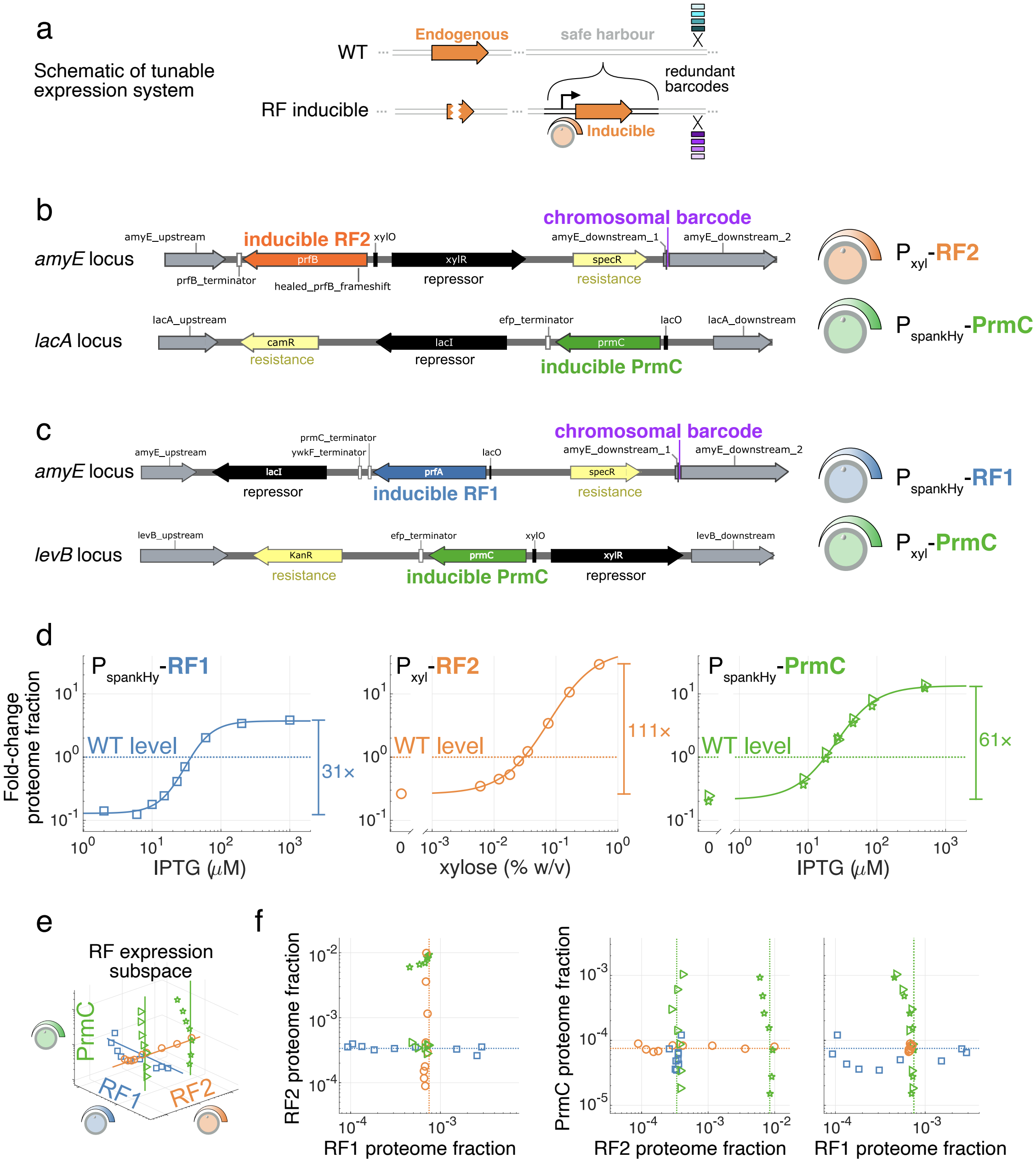

**Fig. S1. Details of RF-inducible expression constructs. Related to Fig. 1 and 2. (a)** Schematic tunable expression system. Inducible constructs are added at safe harbor loci, together with a barcode for competition experiments. The endogenous gene copy is then deleted in a scarless fashion. **(b-c)** Details of loci with tunable expression cassettes for the orthogonally tunable **(b)** RF2 and PrmC strain, and **(c)** RF1 and PrmC strain. Disrupted safe harbor endogenous loci are *amyE*, *lacA*, and *levB*. Control strains were also constructed with blank expression cassettes at these locations (Methods, Fig. S2f-g, S2i-j). Inducible repressors XylR and LacI, respectively responsive to IPTG and xylose are shown in black. Resistance cassettes are shown in yellow. The location of the 8-nt chromosomal barcode in one arm of the *amyE* homology region is shown in purple. **(d)** Fold-change in RF levels as a function of inducers, with fitted Hill curve serve as a guide to the eye. Endogenous expression is shown as the horizontal dashed line (fold-change of 1) and the full attainable dynamic range indicated on the right. See Methods for calibration to proteome fraction. **(e)** 3D RF expression space for all phenotypically profiled conditions (shown separately in Fig. 2a-d) and **(f)** orthogonal projection in 2D subspaces. Dashed lines mark endogenous expression levels.

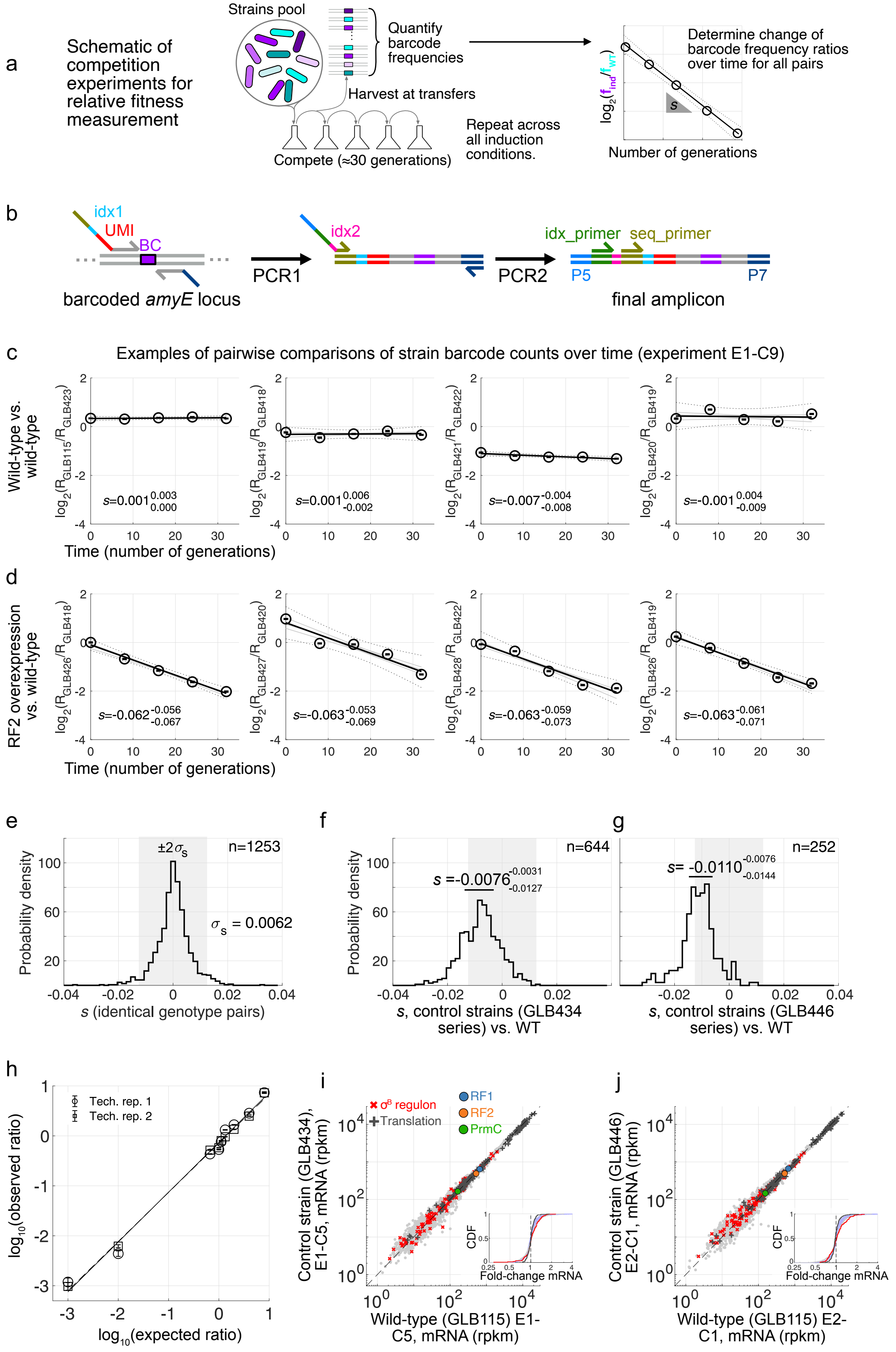

**Fig. S2. Details of relative growth rate measurement. Related to Fig. 2.** Legend on next page.

**Fig. S2. Details of relative growth rate measurement. Related to Fig. 2. (a)** Schematic of competition experiments. Pools of barcoded strains are competed for ≈30 generations with 5 samplings, barcode frequencies are quantified, and changes in barcode frequencies over time determined. Relative growth rate is $\lambda_{\text{inducible}}/\lambda_{\text{WT}} = 1+s$, where $s$ is the slope of the $\log_2$ barcode ratio vs. time (number of generations). This process was performed for all induction conditions shown in Fig. 2a-d. **(b)** Schematic the barcode readout procedure, carried out in two PCR steps from genomic DNA extracted from pools of competing strains, with UMI and first index added at the first PCR, and a second index added at the second PCR. Details of the final amplicon for barcode readout is in Supplementary Data 6. **(c-d)** Examples of barcode frequency ratios over time for isogenic strain pairs, **(c)** wild-type vs. wild-type, and **(d)** RF2 overexpression vs. wild-type. Representative strain pairs from experiment E1-C9 are shown. Inferred $s:=\lambda_{\text{inducible}}/\lambda_{\text{WT}}-1$ from the linear fit (black line) is shown on the graph. Range of slopes $s_{min}$ to $s_{max}$ of subsampled bootstraps (gray lines) is reported as $s_{s_{min}}^{s_{max}}$. **(e)** Distribution of measured $s$ for pairs of strains with identical genotype apart from barcode across all experiments (n=1253 comparisons, e.g., 4/1253 experimentally determined $s$ are shown in panels (c)). The shaded gray area corresponds to the $\pm 2\sigma_s = \pm 1.2\%$ shown in Fig. 2e-h. **(f-g)** Measured fitness difference to wild-type for strains with blank expression cassettes. **(f)** Blank $P_{xyl}$ at *amyE* & blank $P_{spankHy}$ at *lacA*, strains GLB434-437, n = 644 comparisons. **(g)** Blank $P_{spankHy}$ at *amyE* & blank $P_{xyl}$ at *levB*, strains GLB446-449, n=252 comparisons. Both control strain series show minimal effect of ectopic insertions on cell fitness. $s$ values displayed correspond to median with 25th and 75th percentile of values across isogenic pairs. **(h)** Hand mixing experiment with two strains with different barcodes, showing accurate (slopes 1.00 and 0.98 for observed vs. expected ratio of barcodes from two technical replicates) readout of cell frequencies in pool over nearly 4 orders of magnitude. Median difference between readout from two technical replicates is 20%. **(i-j)** Representative examples of mRNA level (RNA-seq) comparison between wild-type and control strains with blank expression cassette insertions: **(i)** GLB434 vs. wild-type (experiment E1-C5), and **(j)** GLB446 vs. wild-type (experiment E2-C1). Cumulative distributions of fold-changes are shown as insets as in Fig. 3a-b.

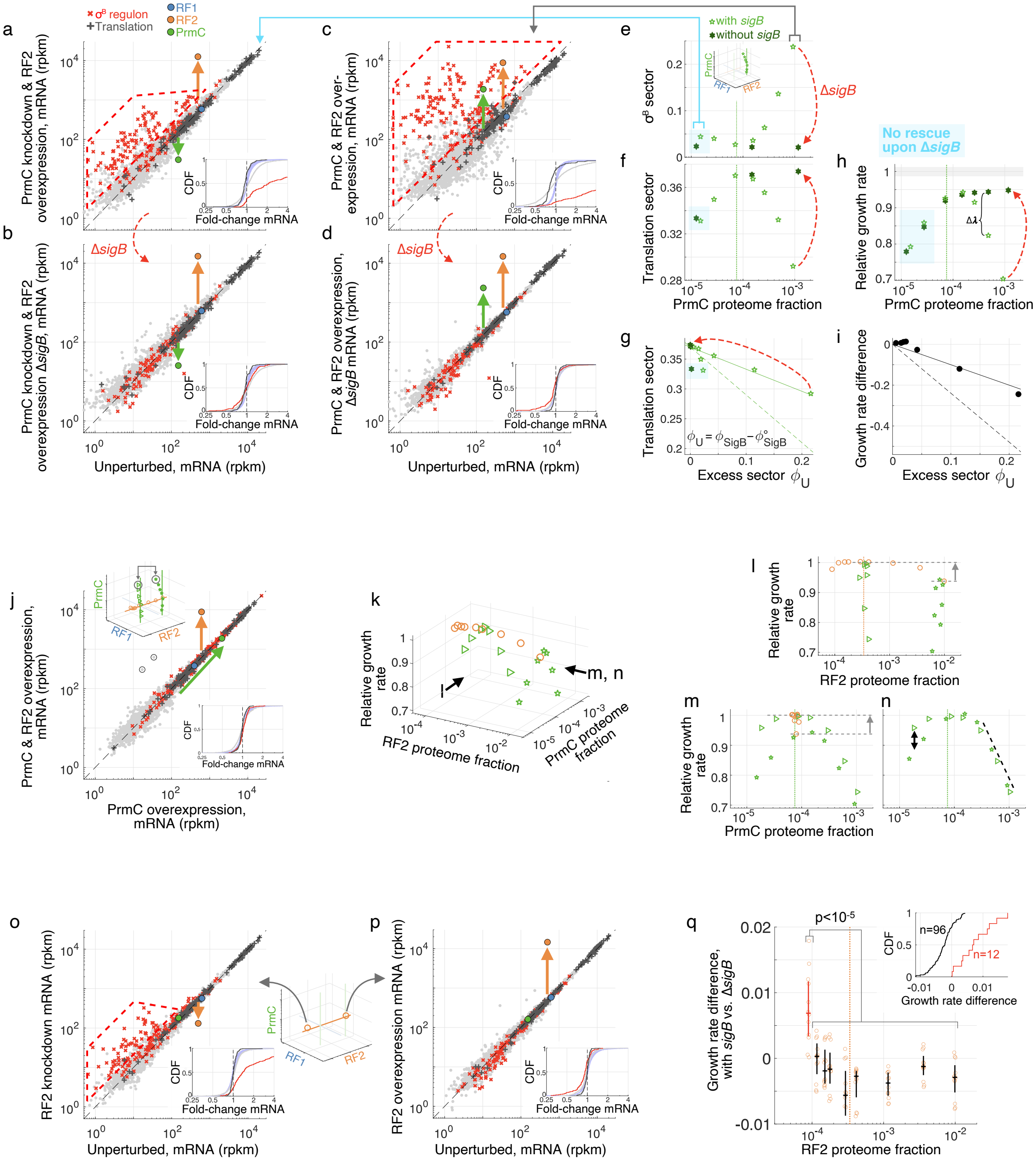

**Fig. S3. Interplay between expression of RF2 and PrmC, fitness and $\sigma^B$ regulon activation. Related to Fig. 3 and 4.** Legend on next page.

**Fig. S3. Interplay between expression of RF2 and PrmC, fitness and $\boldsymbol{\sigma}^B$ regulon activation. Related to Fig. 3 and 4.** Panels **(a-i)** analogous to Fig. 3, but for varying PrmC levels in conjunction with RF2 overexpression (conditions shown in Fig. 2d). In panels **(e-h)**, open light green pentagrams correspond to cells with *sigB*, and filled dark green hexagrams to cells without *sigB* (deletion). Blue shadings mark the region of the expression space for which the growth defect is not rescued by *sigB* deletion, indicating a different underlying cause for the decrease in translation sector. **(j)** Comparison of expression at maximal PrmC expression for endogenous and overexpressed RF2 levels (respective comparisons to unperturbed conditions in Fig. 3a and S3c), showing highly reproducible $\boldsymbol{\sigma}^B$ induction independent of RF2 levels (the two outliers marked by black circles are *xylA* and *xylB*, which are responsive to xylose). **(k)** Expression-fitness landscape for RF2 and PrmC, with orthogonal projections showing the **(l)** RF2, and **(m-n)** PrmC directions. Panel **(n)** is the PrmC fitness landscape, with the fitness defect caused by RF2 overexpression defect subtracted out (arrows in panels (l) and (m)). Fitness defect at overexpressed PrmC is independent of RF2 (dashed black line in (n)). Knockdown defect is exacerbated by RF2 overexpression (black arrows in (n)). **(o-p)** Transcriptome under RF2 expression perturbation. **(o)** RF2 knockdown shows modest $\boldsymbol{\sigma}^B$ induction, whereas **(p)** maximal RF2 over-expression displays little expression changes. **(q)** Growth rate difference for strain with inducible RF2, with and without *sigB*. A mild but significant ($p<10^{-5}$, bootstrap subsampling, Methods) improvement in fitness upon *sigB* deletion at lowest RF2 levels is seen. Measured *s* for each of 12 strain pairs, inducible RF2 (GLB426 to GLB429) vs. inducible RF2 without *sigB* (GLB430 to GLB433), are shown for all profiled RF2 levels, with the median and 25$^{th}$ to 75$^{th}$ percentile marked by black lines. Red marks the condition with lowest RF2 level. Inset shows cumulative distribution of relative fitness difference for lowest RF2 level (red) and rest of conditions (black). The fitness rescue upon *sigB* deletion (median increase in fitness $\Delta s$=0.009) is commensurate with the estimated excess $\boldsymbol{\sigma}^B$ regulon proteome fraction ($\Delta\phi_{\mathrm{SigB}}$ = 0.0085) in this condition.

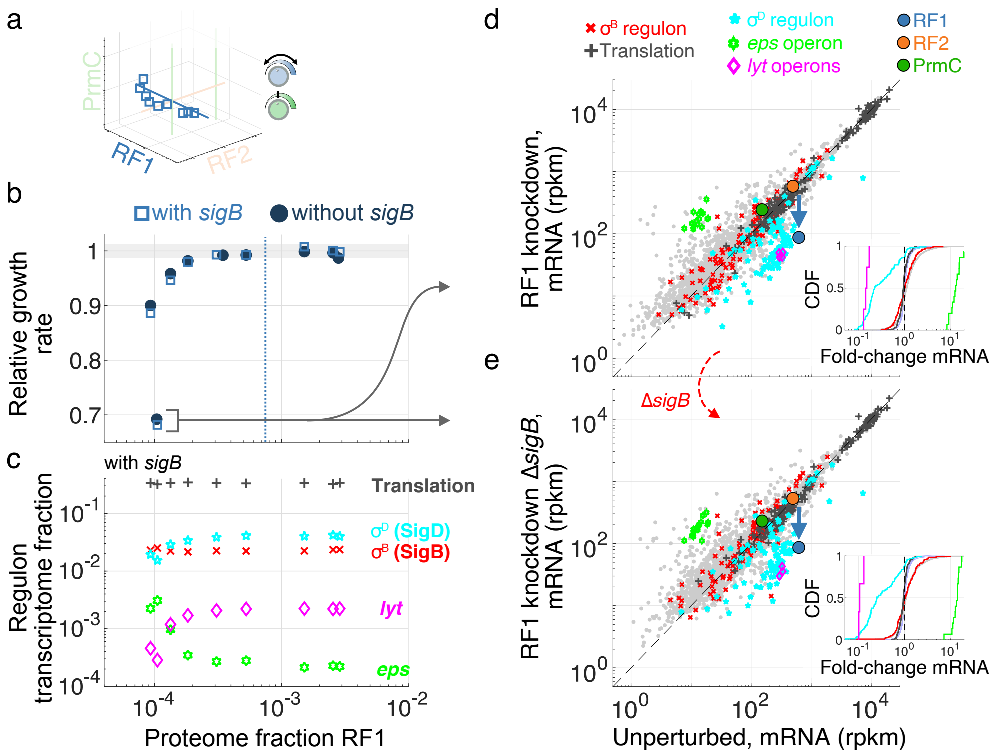

**Fig. S4. Transcriptomics changes upon RF1 knockdown. Related to Fig. 2, 3 and 6. (a)** Profiled RF1 levels in the RF expression subspace (reproduction of Fig. 2a). **(b)** Fitness defect upon modulation of RF1 level, with (open pale blue squares) and without *sigB* (filled dark blue circles), showing no strong influence of $\boldsymbol{\sigma}^{B}$. **(c)** Quantification of various regulons' transcriptome fraction as a function of RF1 levels (shown for strain with *sigB*). The large fitness decrease upon RF1 knockdown coincides with decrease in motility (SigD, cyan) regulon and autolysin operon (magenta), and increase in biofilm matrix *eps* (light green) genes production. **(d-e)** mRNA levels (rpkm, genes with >5 reads mapped shown) comparison to unperturbed for maximal RF1 knockdown, with regulon members colored following (c), for strains **(d)** with or **(e)** without *sigB*. Median fold-change are larger or equal to 5 for SigD, *eps*, and *lyt* genes, independently from $\boldsymbol{\sigma}^{B}$. Cumulative distributions of fold-change for highlighted regulons (rest of genes in gray, all-to-all for across unperturbed replicates in pale blue) are shown as insets as in Fig. 3a.

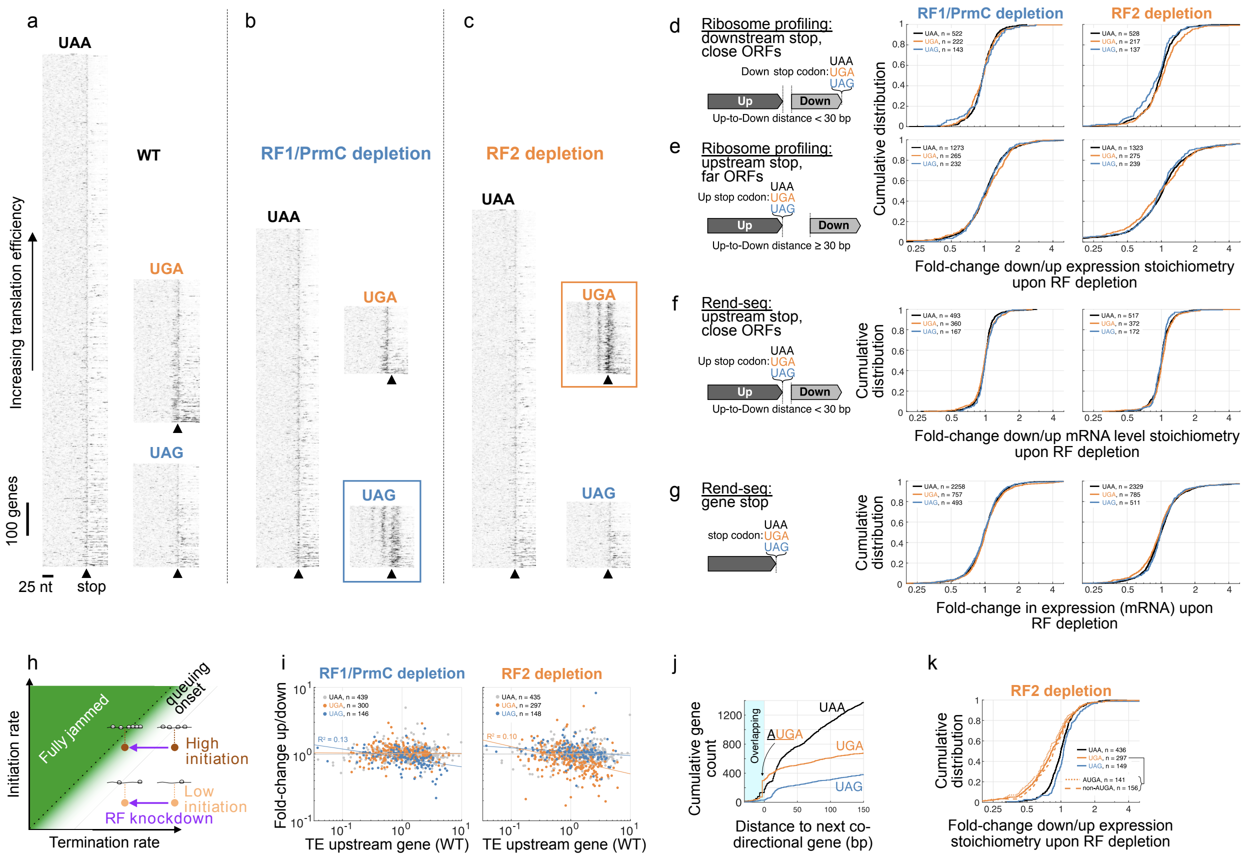

**Fig. S5. Details on translational profiling following acute RF depletion. Related to Fig. 4. (a-c)** Heatmaps of all data used to generate metagene ribosome queuing plot shown in Fig. 4a. for **(a)** wild-type, **(b)** RF1/PrmC CRISPRi depletion, and **(c)** RF2 CRISPRi depletion. Genes are separated by stop codon. Each horizontal line represents a gene, and gene-normalized ribosome footprint density (center-mapped) are shown as gray scale, horizontally aligned by the position of the stop codon (5' to 3' left to right). Genes are organized in increasing order of translation efficiency moving up (TE). Queues upstream of stop codons with perturbed RFs can be seen (colored boxes in (b) and (c)), and are longer for genes with high TE. Scale bars indicate 25 nt, and 100 genes. Caret ▲ marks the position of stop codons. **(d)** Control analysis of changes in expression stoichiometry under RF depletion for co-directional genes within 30 bp, but stratified by the stop codon of the downstream gene (RF1/PrmC: $FC_{UAG}$= 0.99, p=0.39; RF2: $FC_{UGA}$=0.98, p=0.27, p-value from stop codon reshufflings, Methods). **(e)** Similar to (d), but for co-directional genes separated by more than 30 bp, stratifying by the upstream gene (RF1/PrmC: $FC_{UAG}$=1.01, p=0.60; RF2: $FC_{UGA}$=1.04, p=0.96). **(f)** Analysis for expression stoichiometry of gene pairs paralleling Fig. 4d, but with Rend-seq data, showing no effect (RF1/PrmC: $FC_{UAG}$=1.00, p=0.59; RF2: $FC_{UGA}$=1.02, p=0.98). This further suggests that perturbed expression stoichiometry results from changes in translation. **(g)** Distributions of fold-change in mRNA levels between wild-type and RF depletion stratified by stop codon, showing small (≈2%) changes in median for RF-perturbed stop (RF1/PrmC: $FC_{UAG}$=0.98, p=0.06; RF2: $FC_{UGA}$=0.98, p=0.02). **(h)** Schematic illustrating how decreasing termination rate leads to ribosome queues on mRNAs with high translation efficiency (ribosome initiation rate), see Supplementary Discussion. **(i)** Fold-change in expression stoichiometry for gene pairs considered in Fig. 4d (within 30 bp and stratified by upstream stop codon, subset with measured TE shown) as a function of TE. Significant (F-test, $p<0.05$) correlations (increasing translation efficiency leading to more severe effect, $R^2 \approx 0.1$) are seen for genes with stop cognate to the RF perturbation. **(j)** Cumulative number of co-directional gene pairs separated by given distance. Overlap <u>AUG</u>**A** is the most common configuration. **(k)** Same as Fig. 4d, but with UGA pairs split between those with <u>AUG</u>**A** overlap or not, with the overall effect distribution is similar between the two types of overlaps.

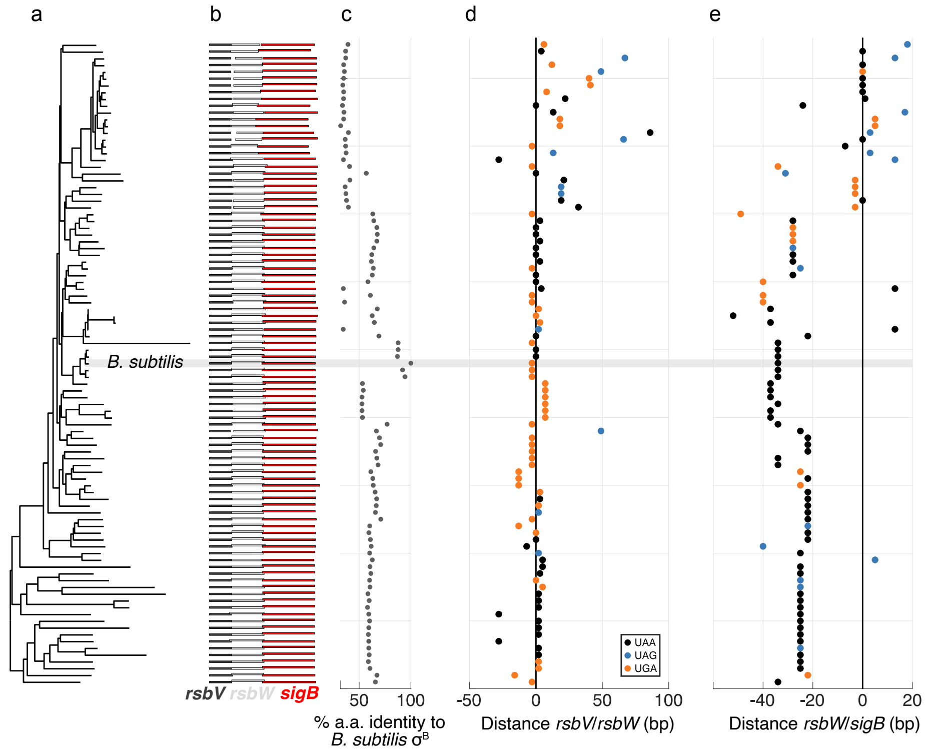

**Fig. S6. $\boldsymbol{\sigma}^{B}$ operon conservation. Related to Fig. 5.** Bioinformatic analysis of $\boldsymbol{\sigma}^{\mathrm{B}}$ operons homologous to that of *B. subtilis* (Methods). **(a)** Phylogenetic tree (neighbor joining from consensus 16S rRNAs) for 95 species with candidate $\boldsymbol{\sigma}^{\mathrm{B}}$ operons (Methods, full list in Supplementary Data 9). **(b)** To-scale miniature representation of gene organization for $\boldsymbol{\sigma}^{\mathrm{B}}$ operon candidates (dark gray: *rsbV* homolog, pale gray: *rsbW* homolog, red: *sigB* homolog). **(c)** Percent identity (amino acid) to *B. subtilis'* $\boldsymbol{\sigma}^{\mathrm{B}}$). **(d-e)** Distance between sequential genes in the operon, colored by stop codon of the upstream gene for **(d)** *rsbV/rsbW* and **(e)** *rsbW/sigB*. 18/95=19% candidate $\boldsymbol{\sigma}^{\mathrm{B}}$ operons have the **<u>AUG</u>A** *rsbV/rsbW* overlap, compared to 226/3042=7% of all co-directional gene pairs in *B. subtilis*. 42/95=44% of the candidate operons have any form of coding sequence overlap (distance < 0 bp) between *rsbV* and *rsbW*, compared to 521/3024=17% across all co-directional gene pairs in *B. subtilis*.
